# Single-cell RNA-seq using UltraMarathonRT expands the known transcriptome

**DOI:** 10.1101/2025.10.06.680646

**Authors:** Chia-Ling Chou, Anastasiya Grinko, Li-Tao Guo, Alexander M. Leipold, Sophie I. Brunets, Teresa Rummel, Florian Erhard, Anna Marie Pyle, Antoine-Emmanuel Saliba

## Abstract

The ability to map messenger RNA (mRNA) molecules from individual cells using next-generation sequencing technologies, known as single-cell RNA-seq (scRNA-seq), is transforming biology by redefining cellular identities with unmatched detail. However, all current protocols depend on copying RNA into complementary DNA with a single reverse transcriptase (RT) derived from murine leukemia virus, which is an RT enzyme known for low processivity and limited ability to unfold complex RNA structures. Here, for the first time, we introduce a group II intron reverse transcriptase, UltraMarathonRT (uMRT), to perform scRNA-seq, a protocol we coined scMarathon-seq. We demonstrate that this enzyme reveals an unexpected transcriptomics landscape by capturing additional genes and other genomic features that conventional RTs miss. We also combined uMRT with metabolic RNA labeling, nucleoside conversion and scRNA-seq to explore genome-wide transcriptome dynamics at the single-cell level. Overall, we establish uMRT as a transformative biotechnological tool for single-cell transcriptomics.

**Summary of changes in Version 2 of the preprint:** The Version #2 of the preprint presents a completely revised scMarathon-seq protocol with enhanced gene detection and gene-body coverage, along with the application of scMarathon-seq to primary cells.

## Introduction

Single-cell (sc)RNA-seq, defined as genome-wide transcriptomics technology at the single-cell level, is significantly transforming cell taxonomy (Tanay and Regev 2017) and uncovering the wide variety of cell states present throughout an organism (Regev et al. 2017). Essentially, the technology’s strength lies in its dimensionality, enabling the measurement of expression levels for thousands of genes simultaneously (Mereu et al. 2020), which allows for a data-driven definition of cell identities (Tanay and Regev 2017). The technology aims to enumerate all RNA molecules in a cell in an unbiased manner, providing a unique identity profile for each cell. Experimentally, scRNA-seq protocols emerged by adapting and improving template-switching-based transcriptomics methods (Ramsköld et al. 2012; Saliba et al. 2014). Such protocols typically follow a three-step process: (i) poly-adenylated messenger RNA molecules are reverse-transcribed into first-strand cDNA; (ii) a primer sequence is added to the cDNA using a template-switching mechanism, allowing the generation of the second strand; and (iii) cDNA is amplified to reach detection sensitivity before sequencing (Saliba et al. 2014). Improvements in the workflow have focused on increasing the number of cells analyzed (Svensson et al. 2018), and enhancing assay sensitivity (Mereu et al. 2020). Additionally, commercial platforms have been developed and deployed to streamline access to the technology (Zheng et al. 2017). However, if a cell can be viewed as a “bag of RNA” (Quake 2021), a natural question arises: do we accurately capture the true picture of RNA molecules in cells?

Central to all scRNA-seq protocols is the ability to reverse transcribe RNA into cDNA using reverse transcriptase (RTs), assuming this step faithfully reflects the RNA census. Published protocols have relied on retroviral RTs derived from a single enzyme originating from Moloney murine leukemia virus (MMLV), which has the power to perform template-switching (Kulpa et al. 1997; Matz et al. 1999). It is produced and marketed in recombinant form with popular names like SuperScript, Maxima, SmartScribe (Zhu et al. 2001; Zucha et al. 2020; Martín-Alonso et al. 2021; Verwilt et al. 2023). However, extensive research has shown that these MMLV variants introduce biases in RNA sequencing due to low processivity and fidelity (Mohr et al. 2013; Guo et al. 2022; Verwilt et al. 2023; Watkins et al. 2026). RT families are not restricted to retroviral RT variants, instead spanning across the tree of life with diverse functions (González-Delgado et al. 2021; Martín-Alonso et al. 2021). Therefore, uncovering the full complexity of the transcriptome requires the use of alternative RTs. Recently, RTs from mobile group II introns have been developed as biotechnology tools for bulk RNA-seq workflows, known as TGIRT, InduroRT, and MarathonRT, exhibiting superior properties compared to MMLV enzymes (Mohr et al. 2013; Zhao et al. 2018). Harnessing these enzymes involves designing the entire RNA-seq process de novo, with the main challenge being to introduce sequence handles for second-strand synthesis and cDNA amplification. Recently, MarathonRT, and its improved version UltraMarathonRT (uMRT), have been shown to perform RNA-seq via a template-switching mechanism, revealing previously unseen transcripts and generating longer cDNA than established MMLV RT (Guo et al. 2024). Given their many advantages, we aimed to incorporate a Group II intron-based RT into the scRNA-seq workflow.

Here, we present scMarathon-seq, a comprehensive plate-based scRNA-seq method using the ultraprocessive enzyme uMRT. We demonstrate that it is suitable for performing efficient scRNA-seq and furthermore uncovers new, unexpected transcriptomic space at the single-cell level compared to the best-in-class method FLASH-seq (Hahaut et al. 2022). Finally, we also establish that our workflow is compatible with scSLAM-seq, encompassing RNA metabolic labeling, nucleotide conversion, and scRNA-seq to measure RNA dynamics of thousands of genes. Overall, our work represents the first integration of ultraprocessive reverse transcriptases into the field of scRNA-seq technologies.

## Results

### scMarathon-seq establishment

In order to adapt UltraMarathonRT (uMRT) for scRNA-seq, we systematically tested a range of reaction conditions for each step of the protocol encompassing cell lysis, mRNA capture, reverse transcription, template-switching and finally, pre-amplification of cDNA libraries (**Figure 1A**). In our first optimization iteration of scMarathon-seq (*BioRxiv* version https://doi.org/10.1101/2025.10.06.680646), we generated cDNA libraries from single K562 cells by introducing betaine in the lysis buffer, Boost reagent in the reverse transcription and reducing the template-switching temperature to 30°C (scMarathon-seq.v1A) and 37°C (scMarathon-seq.v1B). However, as mRNA capture, coverage, and mtRNA levels were suboptimal, we performed a second iteration of optimisations comprising 137 individual experiments, leading to the establishment of scMarathon-seq.v2 (**Figure 1A, Supplementary Table 1**). Given the initial challenges of the protocol, we used cDNA yield and fragment-size distribution as primary read-outs to triage across all conditions. In particular, adjusting reverse transcription to 30°C for 30 min and template-switching to 37°C for 15 min shifted the capture towards longer transcripts, increasing the average library size by 74% with a median length initially of 1,571 bp (start: 460 bp, end: 4,700 bp) jumping to 2,739 bp (start: 480 bp, end: 7,500 bp, **Supplementary Figure 1A**). In addition to the library size, cDNA yield improved, so that PCR cycles could be reduced from 25 to 22. Once we achieved a broad cDNA profile and reproducible yield, we introduced sequencing as a readout, seeking to maximize the number of detected genes. Using 30 pg Universal Human Reference RNA (UHRR), we further tuned the protocol by adapting TSO design, buffer preparation, and Boost and uMRT enzyme concentrations (**Figure 1B**). First, we observed a rise in the number of detected genes from 6,118 (± 359 s.d.) to 9,897 (± 556 s.d.) by increasing Boost concentrations two-fold. While the v2 TSO (RNAConnect) design enhanced gene detection only slightly (10,214 ± 588 s.d.), it lowered TSO concatemer rates from 4.46% (± 2.52 s.d.) to 2.33% (± 1.61 s.d.) and mitochondrial read counts from 19.6% (± 2.19 s.d.) to 17.1% (± 3.21 s.d., **Supplementary Figure 1B**). Using kit-based buffers instead of in-house buffer preparation further increased the number of detected genes to 15,981 (± 359 s.d.) while keeping the other parameters stable. Finally, a maximum of 17,760 (± 1,113 s.d.) genes was detected when increasing uMRT amounts during template switching from 8 to 24 units. Since performing lysis and annealing in one step or in two separate steps did not affect gene detection, we adopted one-step lysis and annealing for easier protocol handling.

**Figure 1:**
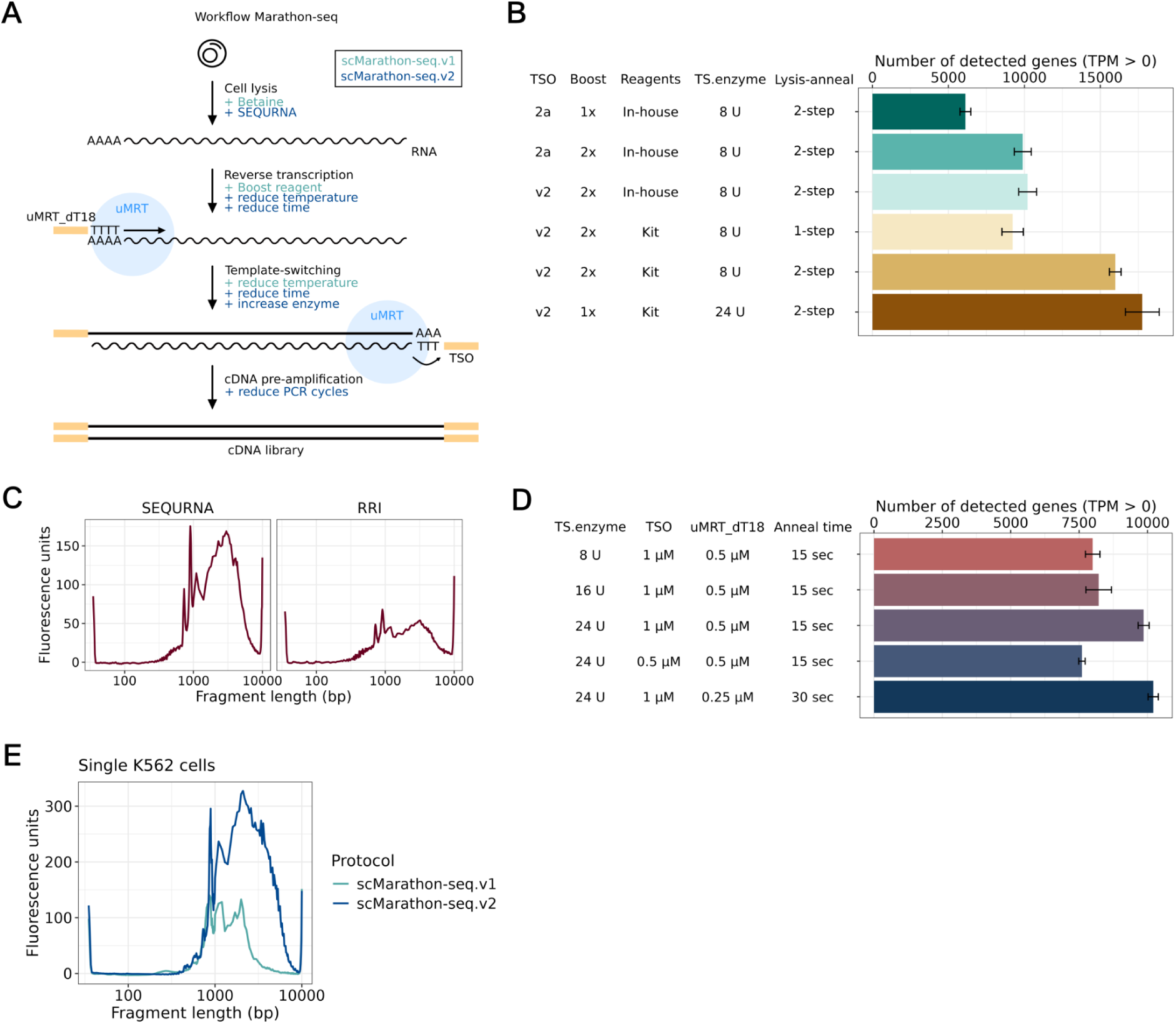
Systematic development and optimization of scMarathon-seq. **(A)** Workflow of scMarathon-seq with highlighted protocol optimization steps for scMarathon-seq.v1 (teal; previous BioRxiv version https://doi.org/10.1101/2025.10.06.680646 version #1) and scMarathon-seq.v2 (dark blue; this version). **(B)** Optimisation of detected genes using 30 pg UHRR as input using different protocol variations. The barplot shows mean genes with standard deviation (n=3) with TPM > 0 for each sample downsampled to 6.9 million reads with varying conditions. Abbreviations: TSO, Template-switching oligo; TS enzyme, amount of uMRT enzyme during template switching in units; Lysis-anneal, lysis and annealing step of the protocol, merged into a single step (1-step) or separated into two steps (2-step). cDNA was amplified with 22 cycles. **(C)** Mean Bioanalyzer traces of cDNA libraries produced from human CD4 cells using SEQURNA inhibitor at 3 U/µL (n = 4) in comparison to recombinant RNase inhibitor (RRI, n = 3). **(D)** Number of detected genes of 10 pg UHRR using different protocol variations (n = 3 for each variation). Abbreviations: TS enzyme, amount of uMRT enzyme during template switching in units; TSO, concentration of template-switching oligo during template switching; uMRT_dT18, concentration of uMRT_dT18 primer during reverse transcription; Anneal time, incubation time of annealing step. cDNA was amplified with 25 cycles. Depicted are genes with TPM > 0 for each sample downsampled to 6.9 million reads. Barplots show the mean and error bars show the standard deviation. **(E)** Mean Bioanalyzer traces of cDNA libraries generated from single K562 cells using scMarathon-seq.v1 (teal, n = 3) and scMarathon.seq.v2 (dark blue, n = 8).

**Supplementary Figure 1, related to Figure 1:**
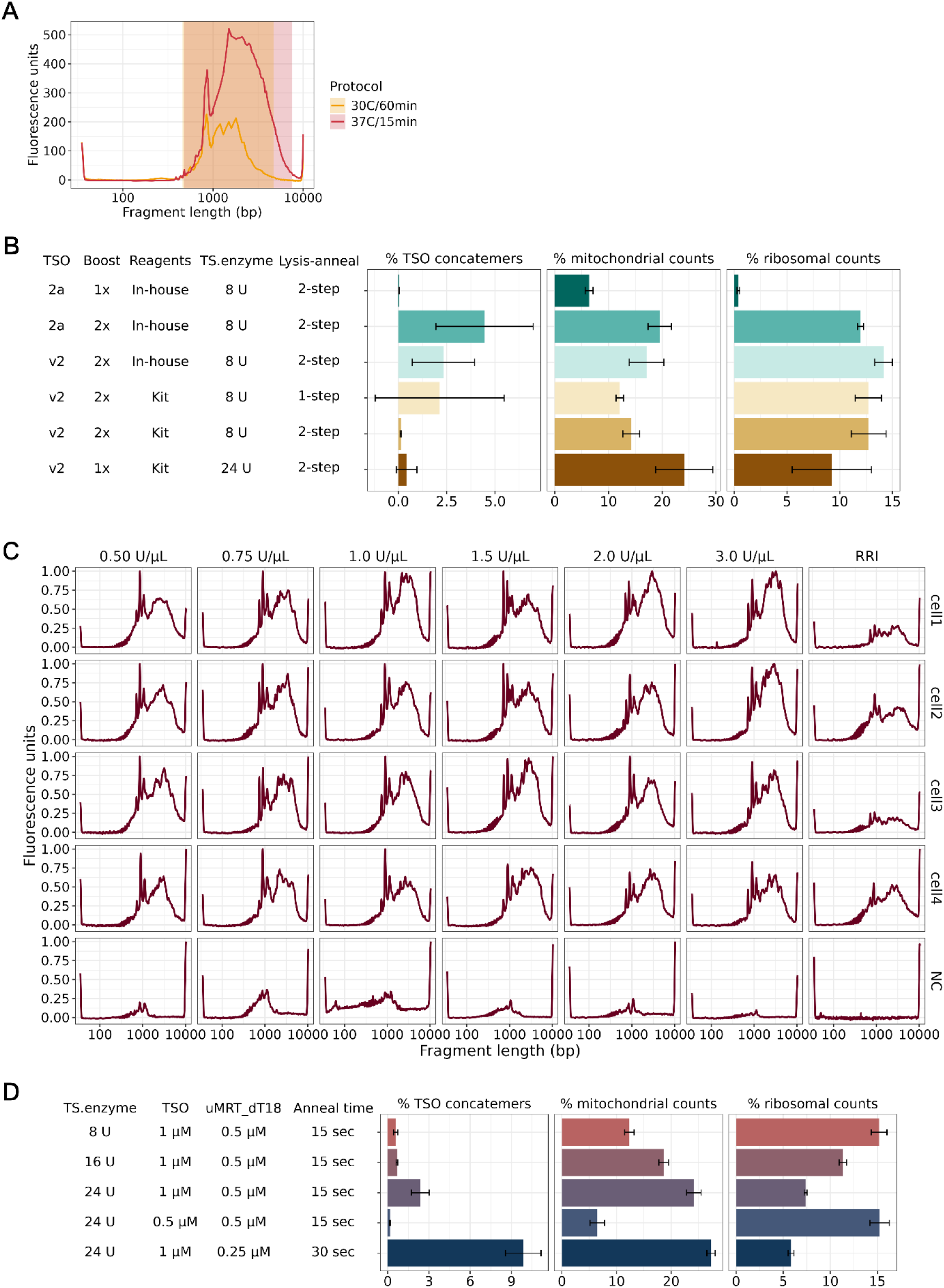
Extended parameters to evaluate scMarathon-seq performance. **(A)** Mean Bioanalyzer traces of cDNA libraries generated from 20-50 pg UHRR with reverse transcription at 42°C for 30 min; template switching at 30°C for 60 min (yellow, n = 3) and from 30 pg UHRR with reverse transcription at 30°C for 30 min; template-switching at 37°C for 15 min (red, n = 3). Library-spanning regions are depicted as rectangular shadows. **(B)** TSO concatemer rates, percentage mitochondrial counts and percentage ribosomal counts of 30 pg UHRR using different protocol variations of scMarathon-seq (n = 3 for each variation). Abbreviations: TSO, Template-switching oligo; TS enzyme, amount of uMRT enzyme during template switching in units; Lysis-anneal, lysis and annealing step of the protocol, merged into a single step (1-step) or separated into two steps (2-step). cDNA was amplified with 22 cycles. TSO concatemer rates were calculated by dividing the number of total TSO concatemer containing reads by the total raw FASTQ reads for each sample. Mitochondrial and ribosomal counts were mapped with STAR and only reads with a MAPQ score of 255 were included for the analysis. Barplots show the mean and error bars show the standard deviation. **(C)** Bioanalyzer traces of cDNA libraries produced from single-sorted CD4 cells using increasing concentrations of SEQURNA inhibitor (n = 4 for each concentration from 0.5-3 mass units/µL (U/µL)) in comparison to recombinant RNase inhibitor (RRI, Takara Bio). NC, negative control. **(D)** TSO concatemer rates, percentage mitochondrial counts and percentage ribosomal counts of 10 pg UHRR using different protocol variations (n = 3 for each variation). Calculations were performed as in **(B)**. TS enzyme, amount of uMRT enzyme during template switching in units; TSO, concentration of template-switching oligo during template switching; uMRT_dT18, concentration of uMRT_dT18 primer during reverse transcription; Anneal time, incubation time of annealing step. cDNA was amplified with 25 cycles. Barplots show the mean and error bars show the standard deviation.

Having enhanced scMarathon-seq’s gene detection over three-fold in 30 pg UHRR, we wondered whether scMarathon-seq could perform at even lower RNA input, with the perspective of applying the protocol to primary cells. To this end, we first explored the use of a novel synthetic, thermostable RNase inhibitor, SEQURNA, which previously showed improved RNA stability at a lower cost (Noble et al. 2024). At concentrations from 0.5 to 3 mass units/µL (U/µL) in the lysis buffer, SEQURNA repeatedly generated higher quality cDNA libraries from sorted human CD4 cells than the standard protein-based recombinant RNase inhibitor (RRI, **Supplementary Figure 1C**). We selected 3 U/µL as the working concentration as it showed the most robust profiles on the Bioanalyzer (**Figure 1C**) and continued with further optimisation experiments using 10 pg of UHRR (**Figure 1D**). Indeed, we were able to detect 7,997 (± 266 s.d.) genes at 8 units, 8,217 (± 469 s.d.) genes at 16 units and 9,860 (± 201 s.d.) genes at 24 units uMRT at 10 pg UHRR, confirming our previous finding that higher uMRT amounts increases template switching efficiency and thus gene capture rates. In parallel, mitochondrial gene capture amplified as well (12.3% ± 0.863 s.d. for 8 units, 18.6% ± 0.875 s.d. for 16 units and 24.1% ± 1.33 s.d. for 24 units uMRT), while TSO concatemer rates remained below 2.5% (0.579% ± 0.150 s.d. for 8 units, 0.671% ± 0.0578 s.d. for 16 units and 2.37% ± 0.651 s.d. for 24 units uMRT) and ribosomal counts even dropping below 8% (15.2% ± 0.8 s.d. for 8 units, 11.3% ± 0.4 s.d. for 16 units and 7.4% ± 0.6 s.d. for 24 units uMRT, **Supplementary Figure 1D**). We tested additional variations in TSO and uMRT_dT18 concentration and annealing time, but lowering the TSO concentration from 1 µM to 0.5 µM only dropped gene capture to 7,603 (± 118 s.d.). Although decreasing uMRT_dT18 concentrations from 0.5 µM to 0.25 µM and extending annealing time from 15 seconds to 30 seconds slightly increased gene numbers by 3.6% (10,211 ± 187 s.d.) and mitochondrial counts by 13.3% (27.3% ± 0.764 s.d.), the TSO concatemer rate raised over three-fold from 2.37% (± 0.651 s.d.) to 9.86% (± 1.29 s.d.). Therefore, we defined 24 units uMRT in template-switching, 1 µM TSO, 0.5 µM uMRT_dT18 and 15 seconds annealing time as our final protocol parameters, which resulted in a dramatic improvement of the cDNA profile of single K562 cells (**Figure 1E**).

Altogether, through adoption of a new TSO, kit reagents, and an increase to 24 units of uMRT for template-switching, scMarathon-seq.v2 shows a three-fold improvement in gene capture, while keeping TSO concatemer rates, mitochondrial and ribosomal counts limited. In combination with SEQURNA, scMarathon-seq.v2 can be applied to RNA input as low as 10 pg, paving the way to profile primary cells.

### Benchmarking scMarathon-seq on the Bulk level

To establish a reference standard for uMRT performance, we extracted 1 ng of total RNA from the ENCODE tier-one cell line K562, as well as the ENCODE tier-two cell line HeLa S3 (also known as CCL-2.2; henceforth HeLa). We compared low input bulk performance of scMarathon-seq to the FLASH-seq (FS) protocol, which is currently one of the top-performer plate-based scRNA-seq protocols in terms of number of genes detected (Hahaut et al. 2022). Since our protocol relies on template switching, we first investigated its impact on our method. While template-switching methods are intended to generate full-length amplicons between primers, uncontrolled template switching can occur, producing TSO concatemers that inflate sequencing libraries (Kapteyn et al. 2010). We found that TSO concatemer rates in scMarathon-seq remained at very low level in the range of 1% to 3% for both cell lines (**Figure 2A**). After read trimming and quality filtering, the majority of high-quality reads mapped uniquely to the GENCODE v47 human reference set with an average 86.3% for HeLa and 83.4% for K562 in uMRT, and 94.4% for HeLa and 92.9% for K562 in FS (**Figure 2B**). scMarathon-seq showed a higher fraction of multimapping reads (12.8% ± 0.32 for HeLa and 15.9% ± 4.0 for K562) than FS (4.48% ± 0.262 and 5.80% ± 0.32, respectively). In contrast, unmapped reads remained low, with less than 1% for both protocols. High intra-replicate correlation was observed for both uMRT (HeLa r = 0.96; K562 r = 0.95) and FS (r = 0.97 for both cell types) (**Supplementary Fig. 2A**). To evaluate sensitivity of both protocols, we first compared their performance on quantifying the molecular abundance of ERCC spike-ins (Baker et al. 2005). Both protocols showed strong linearity across the full dynamic range of ERCC concentrations (**Supplementary Fig. 2B**). We next compared the number of detected genes (TPM > 0) at a downsampled sequencing depth of 7.4 million reads to account for differences in sequencing depth. In HeLa cells, scMarathon-seq detected 15,246 ± 107 genes on average, slightly fewer than FS (16,300 ± 226; **Figure 2C**). In contrast, scMarathon-seq detected more genes in K562 cells (17,224 ± 68) than FS (16,335 ± 73), representing the highest gene detection across all conditions. We further characterized detected genes across biotypes according to GENCODE v47. The number of detected protein-coding genes was similar across protocols, ranging from 11,503 to 11,847 (**Figure 2D**). Differences were primarily observed in other biotypes (**Supplementary Table 2**). In HeLa cells, scMarathon-seq detected fewer lncRNAs than FS (2,924 ± 55 vs. 3,436 ± 143), whereas in K562 cells it detected more lncRNAs (4,076 ± 121 vs. 3,550 ± 108). scMarathon-seq showed a lower tendency to capture processed pseudogenes in both cell types (HeLa: 488 ± 7.78; K562: 824 ± 103) compared to FS (766 ± 3.54 and 862 ± 63.6, respectively). In contrast, scMarathon-seq captured more transcribed unprocessed pseudogenes (TUP), particularly in K562 cells (63 ± 15.6).

**Figure 2:**
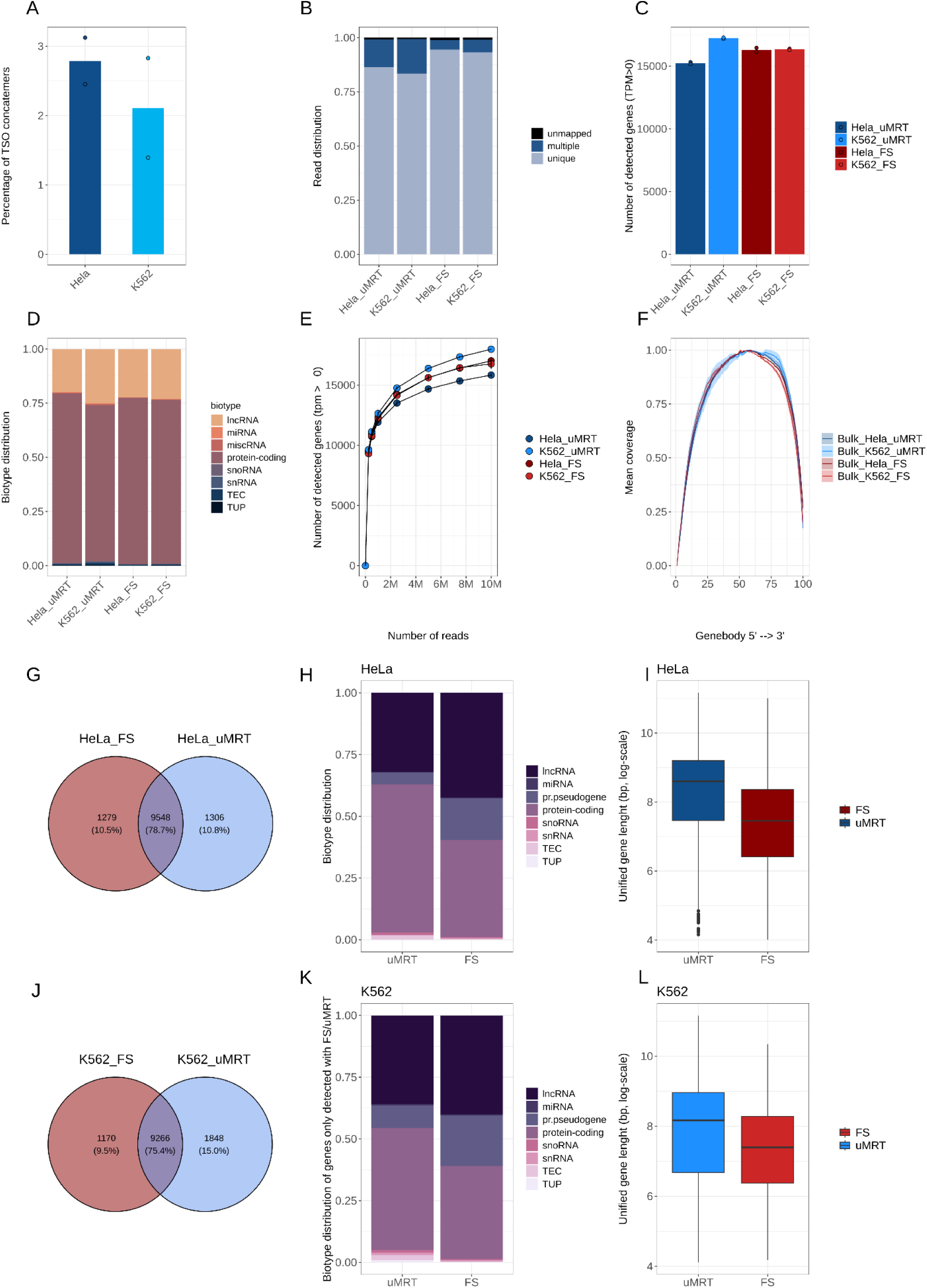
Benchmarking scMarathon-seq on the low input Bulk level. **(A)** uMRT TSO concatemer rates of HeLa (dark blue, n=2) and K562 (light blue, n=2). Barplots indicate mean TSO percentage and points denote single samples. TSO concatemer rates were calculated by dividing the number of total TSO concatemer containing reads by the total raw FASTQ reads for each sample. **(B)** Mapping statistics of uMRT (left) and FS (right) for HeLa and K562 cells. Uniquely mapped reads are denoted in light blue, multiple mapping reads in dark blue and unmapped reads in black. A read is considered to be uniquely mapped with a MAPQ score of 255. Reads were mapped with the STAR aligner. **(C)** Number of detected genes of uMRT HeLa (dark blue, n=2) and K562 (light blue, n=2) and FS HeLa (dark red, n=2) and K562 (light red, n=2). Depicted are genes with TPM > 0 for each sample downsampled to 7.4 million reads. Barplots show the mean between the samples with single sample values shown as points. **(D)** Biotype distribution of detected genes according to GENCODE47. TEC = to be experimentally confirmed, TUP = transcribed unprocessed pseudogene. **(E)** Sequencing saturation curve for uMRT and FS bulk samples. Genes are considered as detected with TPM>0. Downsampling steps were 0.25M, 0.5M, 1M, 2M, 4M, 6M, 8M and 10M. **(F)** Gene body coverage analysis of uMRT and FS HeLa (dark blue and dark red, respectively) and K562 (light blue and light red, respectively) bulk samples. Shown is the full length gene body coverage from 5’ to 3’. **(G)** Gene overlap of HeLa bulk samples between uMRT (blue) and FS (red). For stable expressions, only genes with an expression of TPM > 1 were considered for the comparison. **(H)** Biotype distribution of HeLa genes expressed only in uMRT (left) and only in HeLa (right). Biotypes are named according to GENCODE47. **(I)** Boxplot of gene length of genes captured only by uMRT (dark blue) and only by FS (dark red). Gene length is depicted in bp (log-scale) and taken from the unified gene length per gene of GENCODE47. Boxplots show the median with interquartile range and whiskers indicate the full data range. **(J)** Same as in **(G)** but for K562 cells). **(K)** Same as in (H) but for K562 cells. **(L)** Same as in **(I)** but for K562 cells (light blue = uMRT, light red =FS).

**Supplementary Figure 2:**
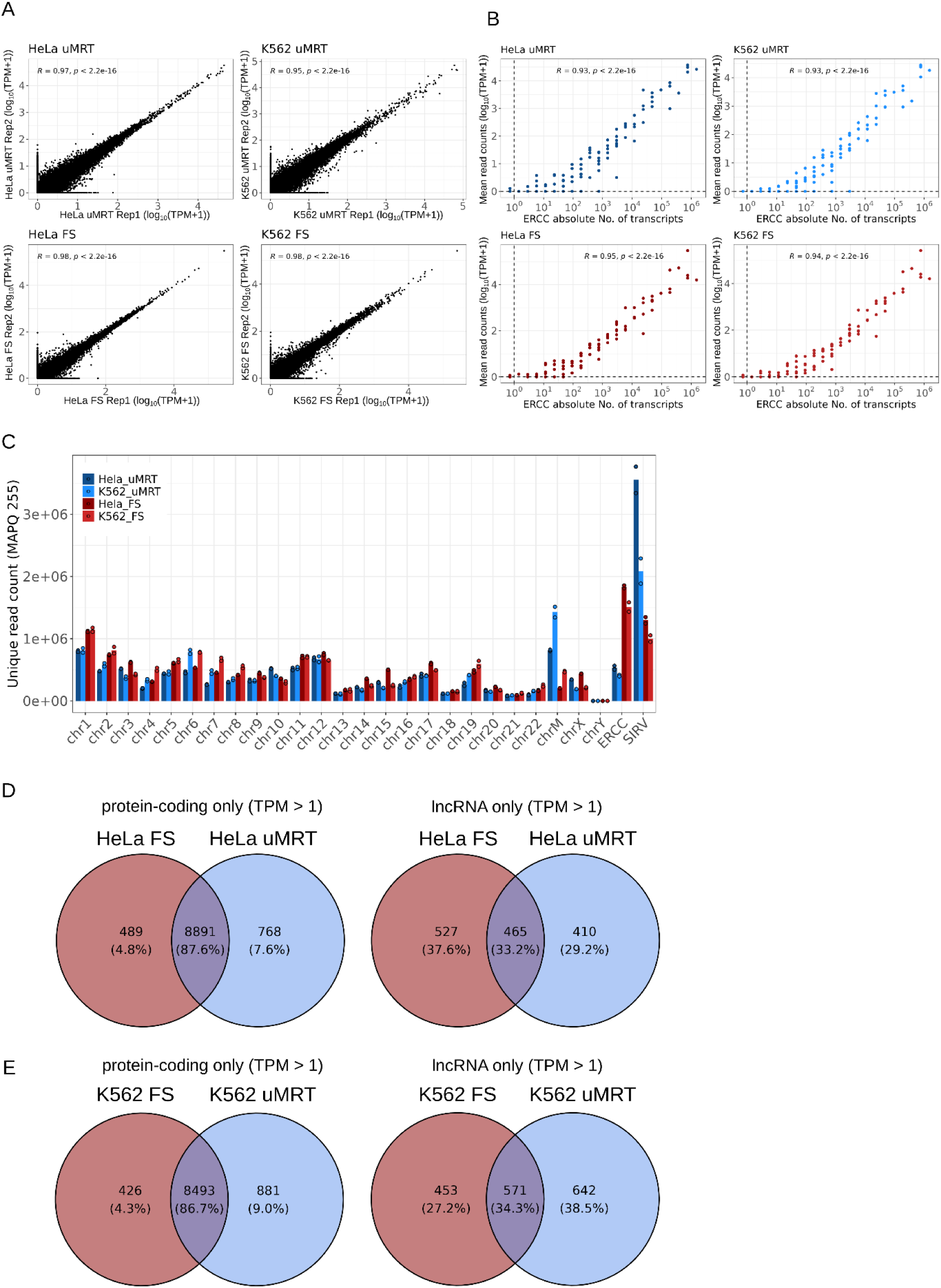
Bulk-level (1 ng) benchmarking of scMarathon-seq. **(A)** Scatter plots of uMRT (upper panel) HeLa bulk samples (left) and K562 bulk samples (right) and FS (lower panel) HeLa bulk samples (left) and K562 bulk samples (right) showing intra-replicate producibility. Pairwise Pearson correlations for expression between samples (uMRT or FS) and replicates were calculated based on log10(TPM + 1) values. **(B)** Sensitivity analysis of uMRT and FS bulk samples using ERCC spike-ins. Absolute number of ERCC molecules is depicted in log10-scale and mean read counts between samples are depicted with log10(+1)-scale. R values were calculated using pairwise Pearson correlation. Upper row (blue shades) shows uMRT samples, lower row (red shades) shows FS samples. **(C)** Distribution of uniquely mapping reads across each human chromosome, including the mitochondrial chromosome and ERCC and SIRV spike-ins. A read is considered uniquely mapped when its MAPQ score is 255. Barplots show the mean number of unique reads with points denoting single samples. **(D)** Intersection of expressed genes between uMRT (blue) and FS (red) focussing on protein-coding (left) and lncRNA genes (right) only for HeLa bulk samples. Only genes with a TPM count > 1 were included in the comparison. **(E)** Same as **(D)** but for K562 cells.

To assess transcript detection completeness, we performed *in silico* downsampling to obtain sequencing saturation curves, simulating sequencing depths of 0.25 to 10 million reads (**Figure 2E**). As expected, the number of detected transcripts increased with sequencing depth. For all samples, plateauing began at around 10 million reads, with K562 samples of the uMRT protocol showing the steepest increase. Moreover, we characterized gene body coverage along the full 5’ to 3’ length of the transcriptome for both scMarathon-seq and FS. Both enzymes showed broadly similar coverage profiles with scMarathon-seq exhibiting slightly more complete coverage toward the 3′ end (**Figure 2F**). Having established key performance metrics, we compared the gene detection space between enzymes. While reads were broadly evenly distributed across all 22 human chromosomes, scMarathon-seq showed higher capture of mitochondrial reads and Lexogen SIRV spike-ins (Paul et al. 2016) (**Supplementary Figure 2C**).

We then evaluated the overlap in detected genes (TPM > 1). In HeLa cells, both protocols shared 78.7% of genes, while approximately > 20% (> 2,500 genes) were uniquely detected by scMarathon-seq or FS (**Figure 2G**). Genes detected exclusively by scMarathon-seq showed a higher proportion of protein-coding genes than those unique to FS. scMarathon-seq also showed enrichment for TEC (“To be experimentally confirmed”) and TUP transcripts, albeit at low abundance (**Figure 2H**). In addition, uMRT-specific genes were longer on average than FS-specific genes (**Figure 2I**). Restricting the analysis to protein-coding genes only increased the overlap to almost 90% (**Supplementary Figure 2D, left**), whereas lncRNAs showed substantially lower concordance (33.2%), suggesting higher variability in lncRNA detection (**Supplementary Figure 2D, right**). These trends were even more pronounced in K562 cells, where 24.5% of genes were uniquely detected by scMarathon-seq (15%) or FS (9.5%) (**Figure 2J**). As in HeLa cells, scMarathon-seq-specific genes were predominantly protein-coding and longer in length (**Figure 2K–L**), with similarly high overlap for protein-coding genes (**Supplementary Figure 2E, left**) and reduced concordance for lncRNAs (**Supplementary Fig. 2E, right**).

Taken together, even at an input as low as 1 ng, scMarathon-seq produces a high-quality bulk transcriptome with low TSO concatenation and gene detection comparable to, or exceeding, FS. Moreover, scMarathon-seq captures a distinct gene space with a distinct biotype distribution profile and enrichment for longer transcripts.

### scMarathon-seq of HeLa and K562

We next assessed scMarathon-seq performance at the single-cell level. We applied scMarathon-seq and FS to FACS (Flow Activated Cell Sorting)-sorted HeLa and K562 cells into lysis buffer-filled plates. We first quantified TSO concatemer rates in single-cell scMarathon-seq libraries. As expected, concatemer rates were higher than in bulk but remained below 7% in both HeLa (6.8% ± 6.8) and K562 cells (6.7% ± 7.1) (**Figure 3A**). The fraction of uniquely mapping reads exceeded 90% for both scMarathon-seq (90.0% ± 2.32 for HeLa; 90.8% ± 1.8 for K562) and FS (93.0% ± 0.9 for HeLa; 94.9% ± 2.32 for K562) **(Supplementary Fig. 3A)**. Using these high-quality reads, we next assessed gene detection in single cells. To account for sequencing depth, we downsampled all libraries to 2 million reads. In addition to our main comparison between optimized scMarathon-seq and FS, we included the previous scMarathon-seq protocol versions (v1A and v1B, *BioRxiv* version https://doi.org/10.1101/2025.10.06.680646) to illustrate methodological improvements, as well as publicly available 10x Genomics datasets from HeLa and K562 cells as droplet-based benchmarks. scMarathon-seq reached a median of 7,053 ± 837 detected genes for HeLa cells and 6,411 ± 1,200 for K562 cells (**Figure 3B**). Strikingly, when comparing this new version of uMRT sequencing conditions to our last scMarathon-seq protocol version, we observed an approximate doubling of the number of detected genes for K562 (4,391 ± 313 for scMarathon-seq v1A and 3,997 ± 386 for uMRT v1B), even at a lower sequencing depth (2 million reads vs. 10 million reads), clearly demonstrating the vast improvements of our new version of the scMarathon-seq protocol. While our new scMarathon-seq protocol detected fewer genes on average than FS for both HeLa cells (8,476 ± 541 vs. 11,417 ± 307, TPM > 0) and K562 cells (8,686 ± 740 vs. 11,113 ± 820), uMRT detected more genes than the standard droplet-based protocols (**Figure 3B**). Correlation between single cells was moderate to high for both protocols and cell types **(Supplementary Figure 3B)**.

**Figure 3:**
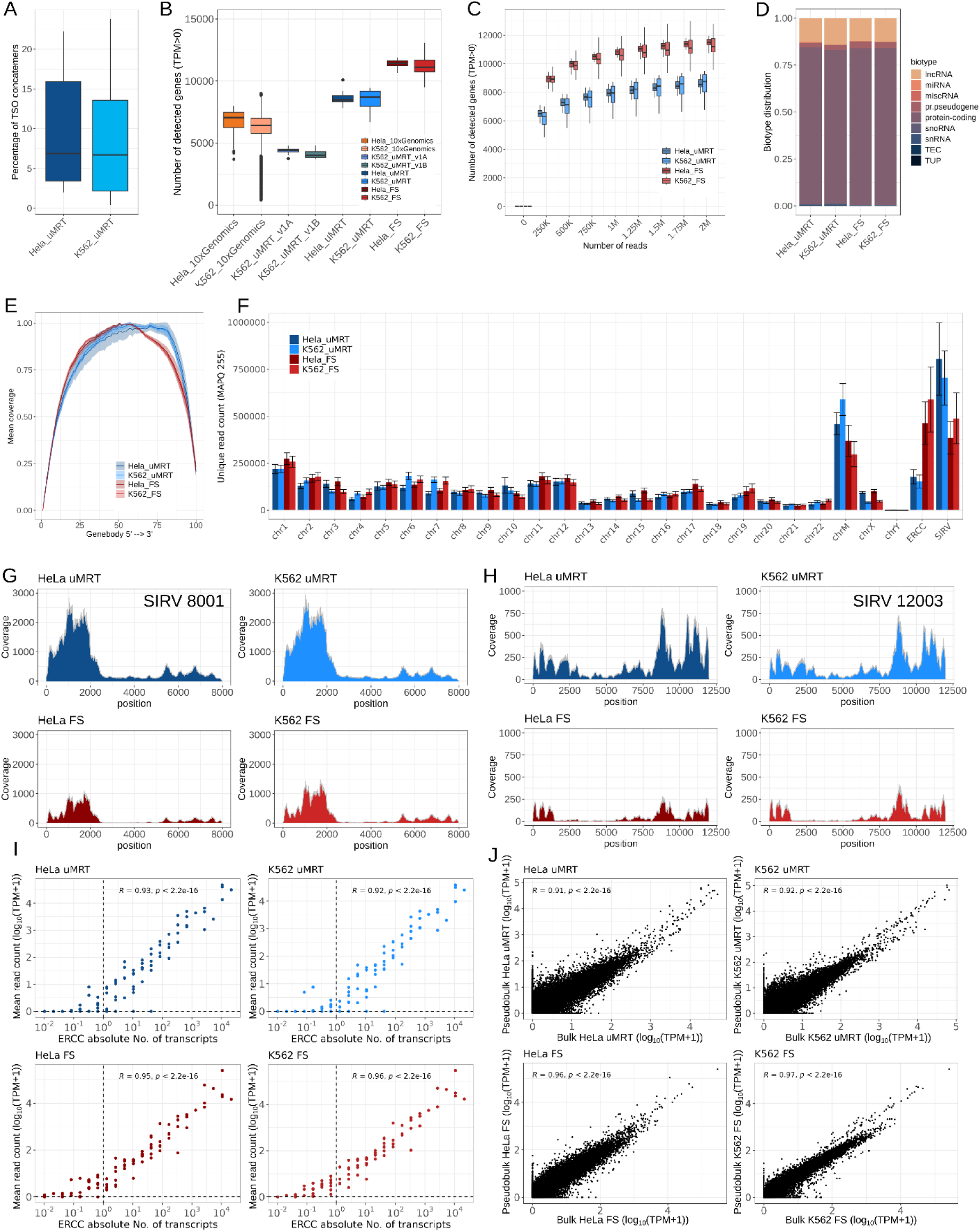
scMarathon-seq applied to profile K562 and HeLa cells. **(A)** scMarathon-seq TSO concatemer rates in HeLa (dark blue, n = 20 cells) and K562 (light blue, n = 20 cells). Boxplots show the median with interquartile range and whiskers indicate the full data range. TSO concatemer rates were calculated by dividing the number of total TSO concatemer containing reads by the total raw FASTQ reads for each sample. **(B)** Comparison of the number of detected genes between multiple single-cell protocols. Shown are the boxplots of detected genes for publicly available 10x Genomics datasets of HeLa cells (dark orange, n=687) and K562 cells (light orange, n=11,542), scMarathon-seq version 1A (pastelblue, n=7), scMarathon-seq version 1B (green, n=7), the current scMarathon-seq version for HeLa cells (dark blue, n=20) and K562 cells (light blue, n=20) and FS for HeLa cells (dark red, n=20) and K562 cells (n=30). Genes are considered as detected with a TPM > 0. **(C)** Boxplots of the sequencing saturation curves for HeLa and K562 cells sequenced with scMarathon-seq and FS. Genes are considered as detected with TPM > 0. Downsampling steps were 0.25M, 0.5M, 0.75M, 1M, 1.25M, 1.5M, 1.75M and 2M. **(D)** Biotype distribution of detected genes according to GENCODE47 for single HeLa and K562 cells of scMarathon-seq (left) and FS (right). TEC = to be experimentally confirmed, TUP = transcribed unprocessed pseudogene. **(E)** Gene body coverage analysis of uMRT and FS HeLa (dark blue and dark red, respectively) and K562 (light blue and light red, respectively) for single-cell samples. Shown is the full length gene body coverage from 5’ to 3’. **(F)** Distribution of uniquely mapping reads across each human chromosome, including the mitochondrial chromosome and ERCC and SIRV spike-ins. A read is considered uniquely mapped when its MAPQ score is 255. Barplots show the mean number of unique reads with error bars denoting the standard deviation. **(G)** Coverage plots of SIRV 8001 (8 kb) of scMarathon-seq (upper panel) HeLa (left, dark blue) and K562 (right, light blue) single cells and FS (lower panel) HeLa (left, dark red) and K562 (right, light red) single cells. Coverage was computed by extracting reads mapping to each genomic position across the transcript and calculating their corresponding depth at that position. Shown is the mean coverage across all cells per sample. **(H)** Same as G but for SIRV 12003 (12 kb). **(I)** Sensitivity analysis of scMarathon-seq and FS single-cell samples using ERCC spike-ins. Absolute number of ERCC molecules is depicted in log10-scale and mean read counts between samples are depicted with log10(+1)-scale. R values were calculated using pairwise Pearson correlation. Upper row (blue shades) shows scMarathon-seq samples, lower row (red shades) shows single-cell FS samples. **(J)** Scatter plots of scMarathon-seq (upper panel) and FS (lower panel) pseudobulk to bulk correlations. Pseudobulks were calculated by aggregating single cell counts per sample. Pairwise Pearson correlations for expression between pseudobulk and their respective bulk samples (scMarathon-seq or FS) were calculated based on log10(TPM + 1) values.

**Supplementary Figure 3:**
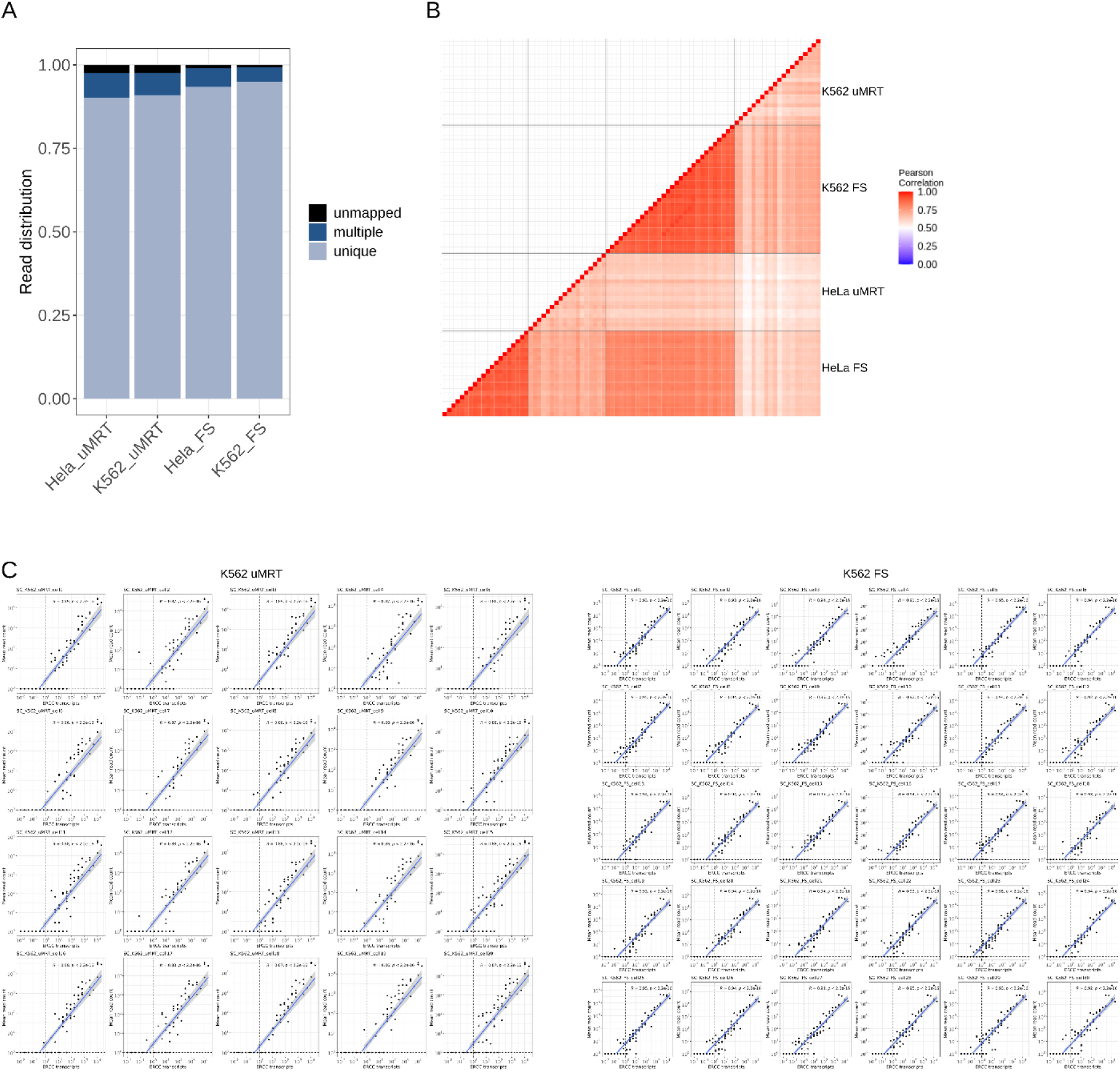
Single-cell RNA-seq metrics of scMarathon-seq. **(A)** Averaged mapping statistics for scMarathon-seq (left) and FS (right) for HeLa and K562 cells. Uniquely mapped reads are denoted in light blue,multiple mapping reads in dark blue and unmapped reads in black. A read is considered to be uniquely mapped with a MAPQ score of 255. Reads were mapped with the STAR aligner. **(B)** Cell-to-cell correlation matrix of uMRT and FS HeLa and K562 cells. Hue corresponds to pairwise Pearson correlation with red indicating high Pearson correlation and blue low Pearson correlation in the range of 0 to 1. For easier visualization, K562 uMRT, K562 FS, HeLa uMRT and HeLa FS samples are divided by grey lines and single cell names are omitted. **(C)** ERCC sensitivity plots for each single K562 cell for scMarathon-seq (left) and FS (right). Absolute number of ERCC molecules is depicted in log10-scale and mean read counts between samples are depicted with log10(+1)-scale. R values were calculated using pairwise Pearson correlation.

As in bulk, we performed *in silico* downsampling to generate sequencing saturation curves for all single-cell libraries, simulating depths from 0.25 to 2 million reads. Gene detection increased with sequencing depth and approached saturation between 1 and 2 million reads indicating that 2 million reads capture most detectable transcripts **(Figure 3C)**. We next examined the biotype distributions of the detected genes. Overall, both protocols showed similar patterns, consistent with bulk results **(Figure 3D)**. To assess differences in detail, we applied a chi-squared test of independence and inspected standardized Pearson residuals to identify category-specific deviations. For protein-coding genes, differences were modest, with a slight decrease in scMarathon-seq K562 cells (Pearson residual = −1.37). However, for lncRNAs, we observed a notable increase for scMarathon-seq in K562 cells (2.9) compared to HeLa cells (-0.24) and a notable decrease in lncRNA capture for both HeLa and K562 cells with FS (-1.6 and -0.6, respectively). As in bulk, scMarathon-seq showed reduced detection of processed pseudogenes (HeLa : −3.3; K562: −2.5) compared to FS (3.5 and 1.4, respectively). In contrast, scMarathon-seq showed enrichment for short non-coding RNAs (miRNA, miscRNA, snoRNA, snRNA) and TEC transcripts **(Supplementary Table 3)**, albeit at low abundance. TUP transcripts were also preferentially detected by scMarathon-seq, particularly in K562 cells (Pearson residual = 2.4). After establishing the captured gene space of scMarathon-seq and FS, we investigated the gene body coverage across the whole 5’ to 3’ length of the transcriptome between both enzymes. Coverage was broadly uniform for both protocols, although uMRT showed a more complete reconstruction with a slight bias to the 3’ end. **(Figure 3E)**. Next, we compared the distribution of reads across each human chromosome and both ERCC and SIRV spike-ins. Read patterns were overall similar, although mitochondrial transcripts and SIRV spike-ins were more strongly represented in scMarathon-seq **(Figure 3F)**.

Given these differences, we examined coverage across individual SIRV transcripts. For that, we specifically focused on long SIRV transcripts, spanning lengths of 4 kb up to 12 kb as a proxy of length complexity. For the 8 kb long SIRV 8001 transcript, we not only demonstrated higher read counts on average, but also higher processivity across the whole transcript from 5’ to 3’ with scMarathon-seq compared to FS (**Figure 3G**). This was also observed for the 12 kb SIRV12003 transcript, where FS showed a stronger drop-off in coverage (**Figure 3H**). Although both transcripts exhibited region-specific coverage biases, uMRT consistently captured reads across regions where FS showed reduced processivity, suggesting increased transcript elongation capability, consistent with previous reports (Guo et al. 2022, 2024). We then assessed scMarathon-seq sensitivity at the single-cell level using ERCC spike-ins. As in bulk, both protocols showed strong linearity across the dynamic range of ERCC input concentrations. scMarathon-seq detected up to five orders of magnitude of ERCC molecules, approximately one log level fewer than FS **(Figure 3I)**. This pattern was consistent across individual cells **(Supplementary Figure 3C)**. Finally, scMarathon-seq pseudobulks of single cells show high correlation to their respective bulk samples (r = 0.91 for HeLa, r = 0.92 for K562), similar to FS (r = 0.96 and r = 0.97, respectively) **(Figure 3J)**.

In summary, scMarathon-seq achieves high performance at the single-cell level. Compared to earlier versions, gene detection is nearly doubled. We observe broadly uniform gene-body coverage with minor biases, high transcript processivity, sensitivity near the lower limit of ERCC detection, and strong correlation to bulk transcriptomes.

### scMarathon-seq captures a different transcriptome than FLASH-seq

Having established uMRT’s high performance at the single-cell level, we next investigated differences in transcriptome capture between scMarathon-seq and FS. We first compared the distribution of genomic elements across all genes. For both enzymes, CDS exons constituted the largest fraction of assigned features **(Figure 4A)**. However, scMarathon-seq showed a lower proportion of CDS exons (54.4% for HeLa and 54.5% for K562) than FS (66.5% and 70.3%, respectively). This difference is largely explained by a higher fraction of intronic reads in scMarathon-seq (19.2% in HeLa and 20.5% in K562) compared to FS (2.9% and 3.2%, respectively). The proportions of 5′ and 3′ UTR exons were similar between both protocols. However, scMarathon-seq showed slightly increased recovery of reads mapping 10 kb downstream of transcription end sites (TES) (0.72% in HeLa and 0.88% in K562) compared to FS (0.14% and 0.19%, respectively), although overall levels in this category were low. A similar trend was observed for regions 10 kb upstream of transcription start sites (TSS), where scMarathon-seq captured 0.24% (HeLa) and 0.41% (K562), compared to 0.06% and 0.08% for FS. Notably, the proportion of unassigned reads was slightly lower for scMarathon-seq (4.7 % for HeLa and K562 cells) than for FS (6.2 % for HeLa cells, 6 % for K562 cells). Next, we examined differences in the gene space between both protocols. Only genes with TPM > 1 and detected in at least 30% of cells were included. In HeLa cells, 71% of genes were shared between scMarathon-seq and FS (**Figure 4B, top**), while the overlap in K562 cells was 70.3% (**Figure 4B, bottom**). Restricting the analysis to protein-coding genes increased overlap to almost 80% in HeLa cells **(Supplementary Fig. 4A, left)** and 78.7% in K562 cells **(Supplementary Fig. 4B, right)**. In contrast, overlap for lncRNAs dropped to less than one-third in HeLa cells **(Supplementary Fig. 4A, right)** and 32.6% in K562 cells **(Supplementary Fig. 4B)**. These results are consistent with bulk data, suggesting that lncRNAs are less consistently recovered across protocols. Although the number of genes uniquely detected by scMarathon-seq was low, we further assessed the biotype composition of these genes using a chi-squared test of independence and standardized residuals. We observed a pronounced enrichment of lncRNAs in scMarathon-seq-only genes (Pearson residual = 7.9 for HeLa cells and 11.2 for K562 cells), compared to FS (−3.5 and −6.4, respectively) (**Figure 4C)**. Additionally, scMarathon-seq-specific genes were overrepresented in the TEC category with Pearson residuals of 7.3 for HeLa cells and 7.6 for K562 cells compared to -3.2 and -3.8 for FS, respectively. While no strong differences were observed for processed pseudogenes, the TUP category was overrepresented in uMRT-specific genes, particularly in K562 cells (Pearson residual = 5.5). Short non-coding RNAs also showed higher residuals in uMRT **(Supplementary Table 4)**. To further characterize global differences between scMarathon-seq and FS, we performed Principal Component Analysis (PCA) excluding mitochondrial and ribosomal reads. PCA revealed two major axes of variation. PC1 (39.1% variance explained) separated HeLa and K562 cells, while PC2 (21.1%) separated scMarathon-seq and FS samples **(Figure 4D)**. Accordingly, samples clustered primarily by cell type (PC1) and secondarily by protocol (PC2), indicating that biological variation exceeded protocol-specific effects. To investigate features underlying this separation, we extracted the 100 and 500 genes with the highest and lowest PC2 loadings. Genes associated with scMarathon-seq (high PC2 loadings) were longer, contained more exons, and had longer intronic regions than FS-associated genes on average **(Figure 4E–G)**. These trends were consistent for both the top 100 and top 500 gene sets, suggesting a robust association between scMarathon-seq and recovery of structurally more complex transcripts. A representative example is *MALAT1*, a long, highly abundant lncRNA spanning approximately 12.8 kb, where we observed distinct coverage patterns across its full length. Although FS recovered a higher number of read mapping to *MALAT1* in HeLa cells, coverage was strongly concentrated at the 5′ end within a 633 bp region spanning chr11:65,499,094–65,499,727 **(Figure 4H)**, where coverage peaked at 8655.85 **(Supplementary Fig. 4C)**. In contrast, scMarathon-seq exhibited more uniform coverage across a larger fraction of the transcript, extending to the 3′ terminus. The same pattern was observed in K562 cells **(Figure 4I)**, where for FS, we observed the same 5’, albeit lower at 6,966 **(Supplementary Fig. 4D)**. We hypothesize that the improved 3′ processivity of uMRT for *MALAT1* is related to non-canonical 3′ end processing of the lncRNA. *MALAT1* lacks a canonical poly(A) tail. Instead, its 3′ end is generated by RNase P cleavage, which simultaneously produces a tRNA-like small RNA (*mascRNA*, ENST00000611300) from the same locus (Wilusz et al. 2008). The mature MALAT1 transcript is stabilized by a short A-rich sequence at its 3′ end, which forms a stable triple-helix structure that protects the RNA from 3′–5′ exonucleolytic degradation (Wilusz et al. 2012). We speculate that uMRT may more effectively traverse this structured 3′ region, enabling more complete transcript coverage. In contrast, FS may show reduced processivity, resulting in read accumulation at the 5’ end.

**Figure 4:**
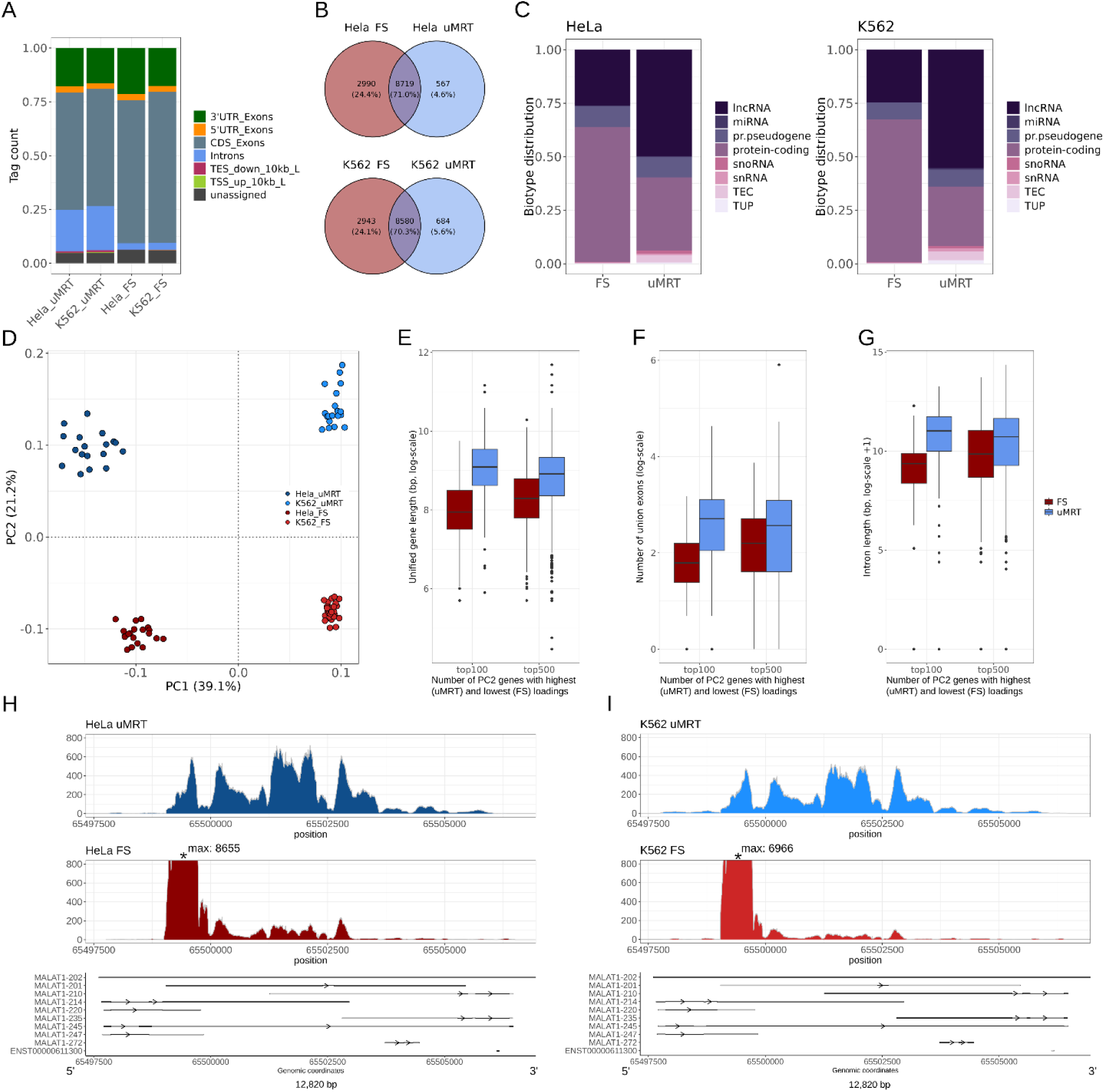
scMarathon-seq captures a different transcriptome than FS. **(A)** Average genomic element distribution between uMRT and FS for HeLa and K562 cells. The distribution was calculated using the RSeQC toolkit based on the GENCODE47 annotation. Only uniquely mapping reads (MAPQ 255) were included in the analysis. A tag corresponds to a read mapping to a certain genomic element. **(B)** Gene overlap of HeLa single-cells (upper panel) and K562 single cells (lower panel) between scMarathon-seq (blue) and fS (red). For stable expressions, only genes with an expression of TPM > 1 expressed in at least 30 % of cells per condition were considered for the comparison. **(C)** Average biotype distribution of HeLa genes expressed only in scMarathon-seq or only in FS in HeLa cells (left) and K562 cells (right). Biotypes are named according to GENCODE47. **(D)** Principal component analysis (PCA) of genes driving variability between HeLa (darker shades) and K562 cells (lighter shades) and uMRT (blue) and FS (red). PCA was calculated on the 2000 most variable genes, using 10 principal components. Mitochondrial and ribosomal genes were excluded from the analysis. **(E)** Boxplot of gene length distribution of PC2 loading genes. Genes with high loading values for PC2 denote uMRT, genes with low loading values for PC2 denote FS. We included both the top 100 and top 500 PC2 separating genes into the analysis. Gene length was taken from the unified gene length of the GENCODE47 annotation. **(F)** Same as in **(E)** but for the number of union exons. The number of union exons was calculated based on the GENCODE47 annotation. **(G)** Same as **(E)** and **(F)** but restricted to intronic length calculated from the GENCODE47 annotation. **(H)** Coverage plots of *MALAT1* (12.8 kb) HeLa single cells of uMRT (upper panel, dark blue) and FS (lower panel, dark red). Coverage was computed by extracting reads mapping to each genomic position across the transcript and calculating their corresponding average depth at that position. Shown is the mean coverage across all cells per sample. Coverage for FS is cut at 800 in order to ensure easy comparability between protocols. The black star indicates the coverage peak of FS. MALAT1 and MALAT1 associated small RNA (mascRNA) were computed from the GENCODE47 annotation. An extracted assortment of MALAT1 isoforms is depicted for simplicity. **(I)** Same as **(H)** but for K562 cells.

**Supplementary Figure 4:**
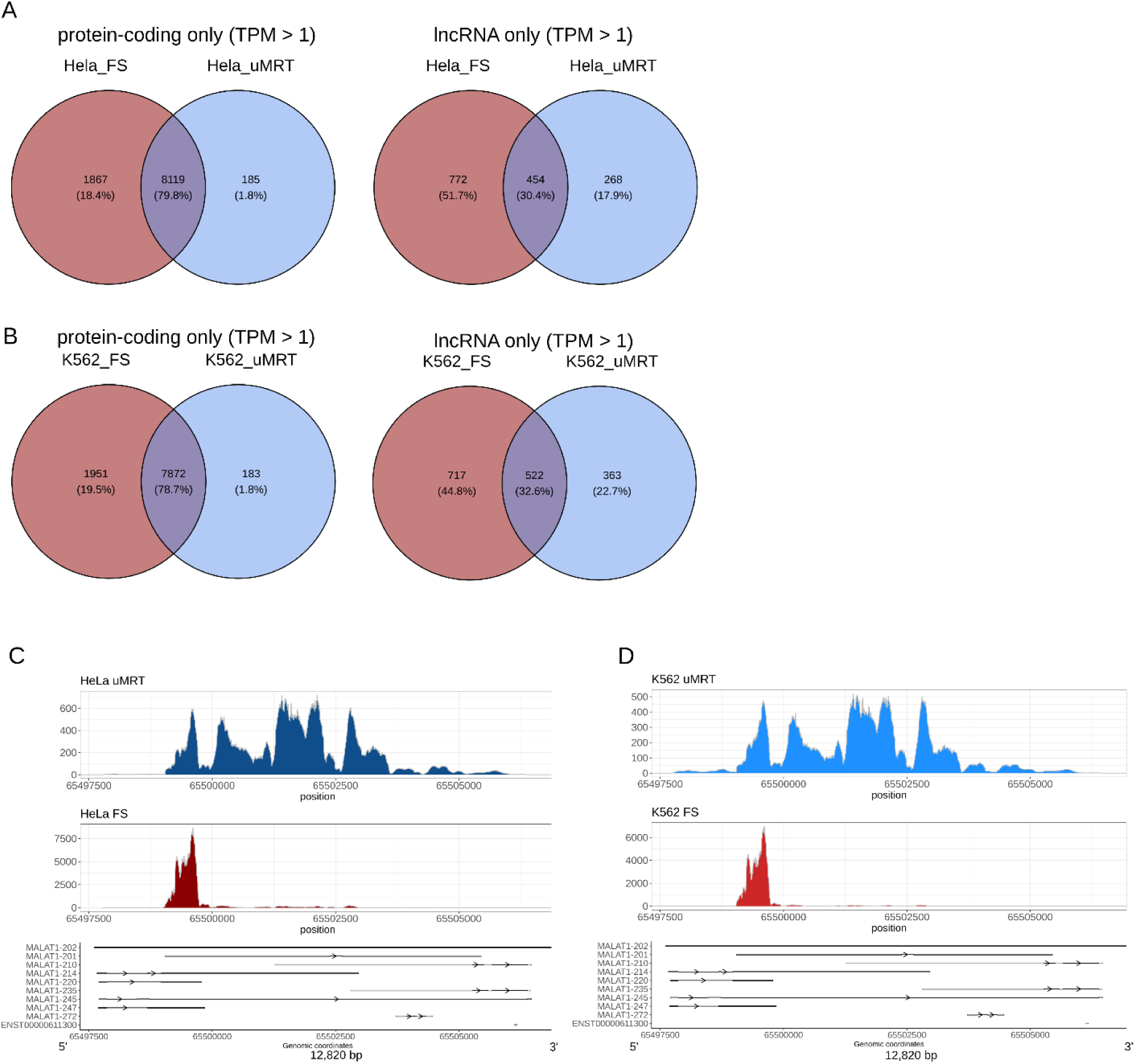
Differences in transcriptome capture between scMarathon-seq and FS. **(A)** Intersection of expressed genes between uMRT (blue) and FS (red) focussing on protein-coding (left) and lncRNA genes (right) only for single cells. Only genes with a TPM count > 1 that were expressed in at least 30 % of cells were included in the comparison. **(B)** Same as **(A)** but for K562 cells. **(C)** Coverage plots of *MALAT1* (12.8 kb) HeLa cells of uMRT (upper panel, dark blue) and FS (lower panel, dark red). Coverage was computed by extracting reads mapping to each genomic position across the transcript and calculating their corresponding average depth at that position. Shown is the mean coverage across all cells per sample. The full range of coverage is shown for FS. *MALAT1* and *MALAT1 associated small RNA* (mascRNA) were computed from the GENCODE47 annotation. An extracted assortment of *MALAT1* isoforms is depicted for simplicity. **(D)** Same as **(C)** but for K562 cells.

In summary, we demonstrate that scMarathon-seq expands the known transcriptome on the single-cell level. scMarathon-seq enriches for longer and structurally more complex genes and shows improved recovery of full-length transcripts. This is underlined by enhanced coverage of *MALAT1* across its full gene body, highlighting the increased processivity of uMRT.

### scMarathon-seq shows a high capture efficiency of the mitochondrial transcriptome

One of the most striking differences we observed between scMarathon-seq and FS was the preferential capture of mitochondrial transcripts. In HeLa cells, 12.8% ± 1.67 of all uniquely mapping reads were assigned to mitochondrial genes in scMarathon-seq, compared to 9.93% ± 2.27 in FS (**Figure 5A**). This effect was more pronounced in K562 cells, where 16.2% ± 2.34 of reads corresponded to mitochondrial transcripts, compared to 8.09% ± 1.87 in FS (**Figure 5A**). Given that elevated mitochondrial reads are often associated with cell stress, lysis artifacts, or differences in cell integrity in single-cell experiments, we additionally quantified mitochondrial read fractions in our bulk samples as a benchmark. Bulk data (**Figure 2**) recapitulated the single-cell trend, with a stronger separation between both enzymes. In HeLa bulk samples, 6.37% of uniquely mapping reads fell within a mitochondrial transcript in scMarathon-seq, compared to 1.46% in FLASH-seq (**Supplementary Figure 5A**). Similarly, in K562 bulk samples, scMarathon-seq recovered 11.6% mitochondrial reads compared to 3.47% for FS. In lieu with that, we next assessed the distribution of reads across single mitochondrial genes. Of the 37 annotated mitochondrial genes (GENCODE47), we focused on the 20 most highly expressed genes. Here, we observed that for FS mitochondrial reads are strongly concentrated on the genes *MT-RNR2*, *MT-ND4L*, and *MT-CO2*, particularly in HeLa cells **(Figure 5B)**. In contrast, scMarathon-seq showed a more even distribution across mitochondrial genes and generally higher read numbers across most transcripts. Even among lower-abundance genes within this set, scMarathon-seq showed higher capture for all genes except *MT-ND3* **(Figure 5B, zoom)**. This trend was amplified in bulk data, where mitochondrial gene expression shifted further in favor of scMarathon-seq. While *MT-RNR2* remained the most highly expressed gene in FLASH-seq for both HeLa and K562, scMarathon-seq exceeded it in TPM as well as for *MT-ND4L* **(Supplementary Figure 5B)**. The only genes with higher expression in FLASH-seq were *MT-CO2* and a subset of low-abundance transcripts, primarily in HeLa bulk samples, but not in K562. These differences were also reflected in coverage across the full mitochondrial genome. In both HeLa and K562 cells, FS showed dominant peaks at *MT-RNR2*, as well as at *MT-CO2* and *MT-ND4L* **(Figure 5C and 5D)**. However, coverage patterns were not directly mirrored between the two protocols. While these peaks were also present in scMarathon-seq, additional transcripts such as *MT-RNR1*, *MT-CO1*, and *MT-CYB* were predominantly captured by scMarathon-seq. This pattern was consistent across both cell types, with higher overall coverage in K562 cells. Additionally, even within individual genes, such as the highly expressed *MT-RNR2* locus, coverage profiles differed markedly between protocols, indicating distinct internal read distributions rather than simple scaling effects. Also, we observed that scMarathon-seq extended coverage toward both the 5′ and 3′ ends of the mitochondrial genome. In FS, coverage typically began at *MT-RNR2*, whereas uMRT also captured *MT-RNR1* at the 5’ end and extended further toward *MT-CYB* at the 3′ end. In bulk samples, these differences were more pronounced, with scMarathon-seq showing substantially higher coverage overall. In particular, K562 bulk scMarathon-seq samples exhibited peaks reaching up to 80,000 reads per base **(Supplementary Fig. 5C and 5D)**. Genes such as *MT-RNR1*, *MT-CO1*, and *MT-ND4* showed particularly strong coverage even at the low input level for the bulk samples. Furthermore, similar as to the single cells, we did not observe mirroring of coverage between scMarathon-seq and FS bulk samples. Indeed, taking the gene *MT-RNR2* again as an example, we continuously demonstrate a dissimilar coverage peak distribution within a single gene, rather than mere differences in magnitude.

**Figure 5:**
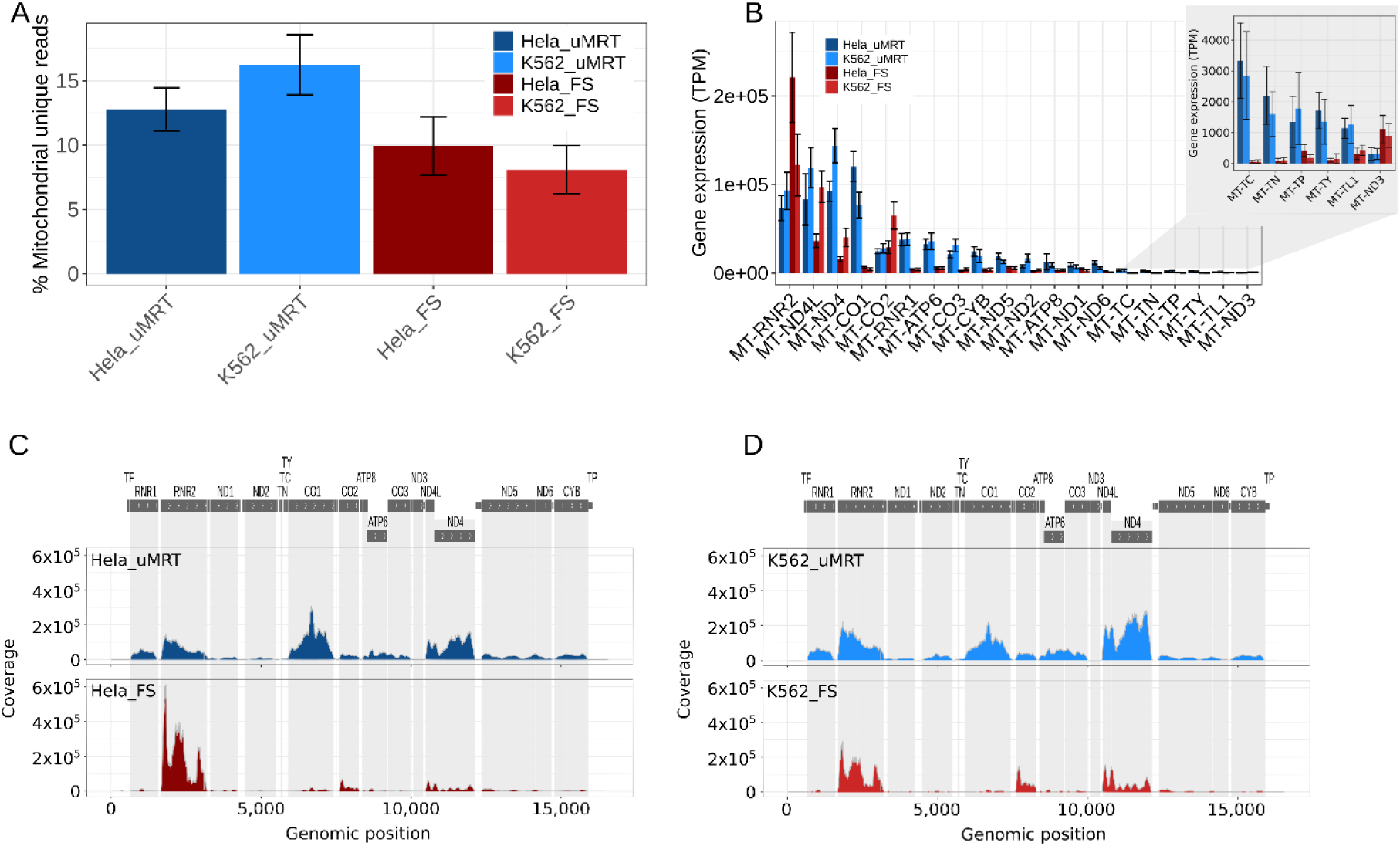
scMarathon-seq captures the mitochondrial transcriptome. **(A)** Percentage of reads falling within mitochondrial transcripts for uMRT HeLa (dark blue) and K562 cells (light blue) and FS HeLa (dark red) and K562 cells (light red). Reads were mapped with STAR and only reads with a MAPQ score of 255 were included for the analysis. Bars depict the mean of all cells with error bars indicating the standard deviation between single cells. **(B)** Gene expression of the top20 most highest expressed mitochondrial genes between scMarathon-seq and FS. Gene expression is shown in TPM and mitochondrial gene annotation was extracted from GENCODE47. The smaller barplot within the grey window highlights the expression of mitochondrial genes expressed at lower ends. Barplots depict mean expression of cells with error bars indicating standard deviation between single cells. **(C)** Coverage plots of the entire mitochondrial genome (16,569 bp) of HeLa cells for uMRT (upper panel, dark blue) and FS (lower panel, dark red). Coverage was computed by extracting reads mapping to each genomic position across the mitochondrial genome and calculating their average depth at that position. Shown is the mean coverage across all cells per sample. Grey boxes on top of the plot depict the annotation of the top 20 highest expressed mitochondrial genes according to GENCODE47 and were created with karyoplotR. **(D)** same as in **(C)** but for K562 cells.

**Supplementary Figure 5:**
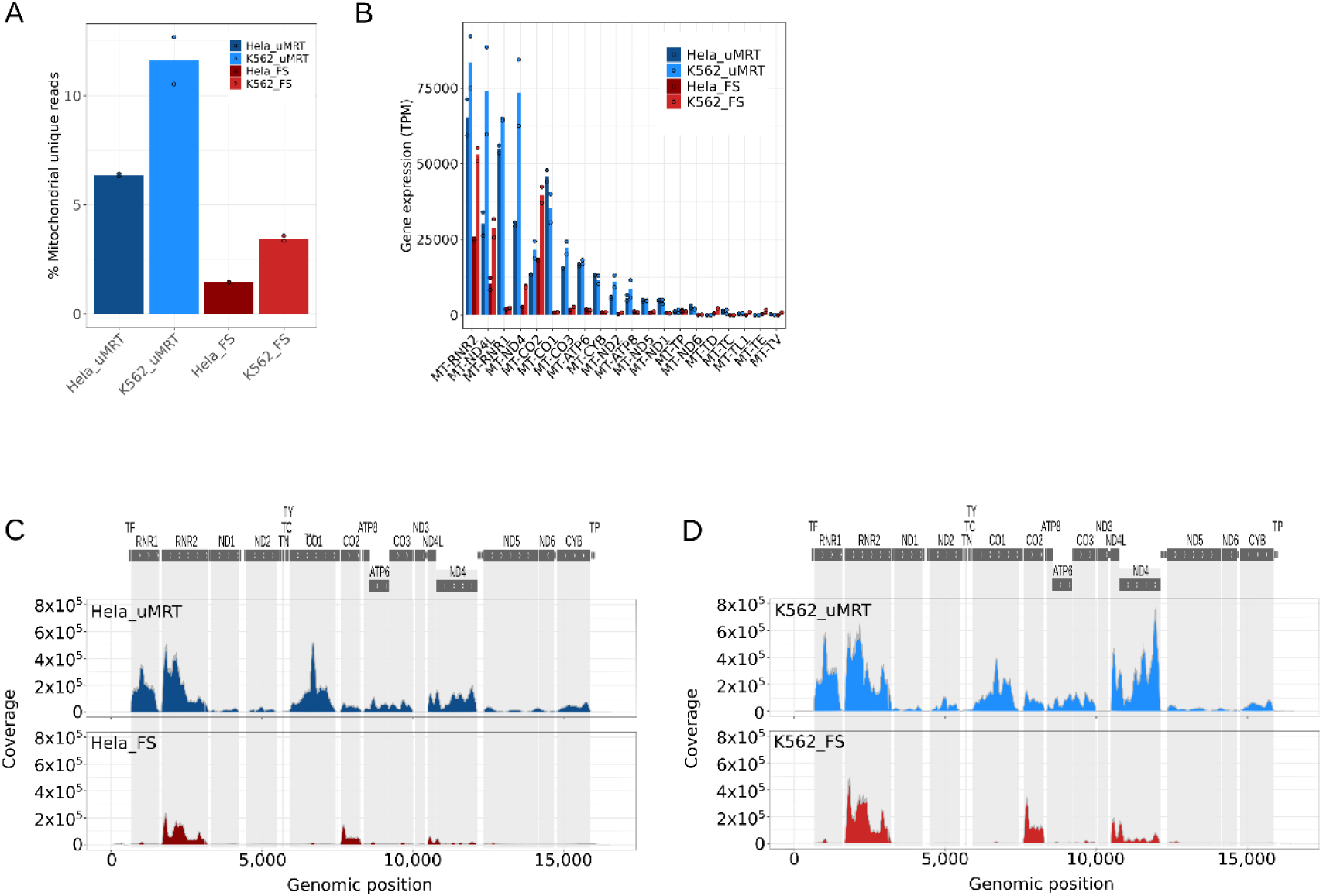
Mitochondrial capture of scMarathon-seq and FS on the bulk level. **(A)** Percentage of reads falling within mitochondrial transcripts for uMRT HeLa (dark blue) and K562 bulk samples (light blue) and FS HeLa (dark red) and K562 (light red) bulk samples. Reads were mapped with STAR and only reads with a MAPQ score of 255 were included for the analysis. Bars depict the mean of all cells with error bars indicating the standard deviation between single cells. **(B)** Gene expression of the most highly expressed mitochondrial genes between scMarathon-seq and FS. Gene expression is shown in TPM and mitochondrial gene annotation was extracted from GENCODE47. Barplots depict mean expression of replicates with points denoting single samples. **(C)** Coverage plots of the entire mitochondrial genome (16,569 bp) of HeLa bulk samples for uMRT (upper panel, dark blue) and FS (lower panel, dark red). Coverage was computed by extracting reads mapping to each genomic position across the mitochondrial genome and calculating their average depth at that position. Shown is the mean coverage across all cells per sample. Grey boxes on top of the plot depict the annotation of the top 20 highest expressed mitochondrial genes according to GENCODE47 and were created with karyoplotR. **(D)** same as in **(C)** but for K562 bulk samples.

Taken together, scMarathon-seq not only shows an increased affinity for capturing mitochondrial transcripts, it also captures these transcripts with a more even distribution across the whole mitochondrial genome. These results suggest that uMRT may be a powerful tool for to elucidate the mitochondrial chromosome.

### scMarathon-seq applied to uncover the transcriptome of PBMC

Finally, we wanted to extend scMarathon-seq to profile peripheral blood mononuclear cells (PBMCs). We specifically sorted for monocytes (CD14), B cells (CD19) and T cells (CD3) cells and prepared them using both scMarathon-seq and FS. In contrast to the HeLa and K562 single cells, we observed TSO concatemer percentages of around 40 % for each PBMC cell type (43.44 % ± 8.79 for CD14, 39.33 % ± 9.1 for CD19, 43.01 % ± 8.79 for CD3) (**Supplementary Figure 6A**). This was reflected in the lower fraction of uniquely mapping reads obtained with scMarathon-seq (63.21 % ± 8.94 for CD14, 66.82 % ± 8.74 for CD19, 60.92 ± 11.05) compared with FS (90.78 % ±1.59, 86.33 ± 7.15 and 87.24 % ± 2.3, respectively) (**Supplementary Figure 6B**) as well as lower gene body coverage (**Supplementary Figure 6C**). Despite these technical limitations, scMarathon-seq was still able to clearly distinguish CD14, CD19 and CD3 cells downsampled to 1 million reads to account for sequencing depth variability. PCA resolved three main clusters along the most variable components in addition with a clear distinction between scMarathon-seq and FS (**Figure 6A**). Furthermore, PC3 captured the technical variability between scMarathon-seq and FS (**Supplementary Figure 6D**). Together, this indicates that biological identity remains the dominant source of variability, rather than the sequencing protocol. We next evaluated whether scMarathon-seq can recapitulate biological signatures. We calculated module scores (see Methods) for each cell type using genes representative of their respective biological programs and projected these scores onto the PCA clusters (**Figure 6B**). Both reverse transcriptases recovered the expected signatures. Moreover, although the space for gene detection was limited in scMarathon-seq due to the substantial number of TSO concatemers, uMRT still detected a median of nearly 2,000 genes across all cell types (**Figure 6C**). For both RTs, CD14 showed the highest number of detected genes (1,905 ± 460 for uMRT, 4,803 ± 739 for FS), followed by CD3 cells (1,811 ± 544 and 3,460 ± 513, respectively) and CD19 cells (1,696 ± 870 and 3,127 ± 709, respectively). To place the number of detected genes into a broader context, we compared our data to publicly available PBMC datasets. We have choosen the benchmarking study by Mereu et al. (Mereu et al. 2020) compared the performance of 13 different single-cell sequencing methods on human peripheral blood. We leveraged the breadth of this dataset to position scMarathon-seq on a global performance scale. Overall, our reanalysis of the data reproduced the results reported by Mereu et al. As expected, plate-based methods showed the highest number of detected genes, with CEL-Seq2 and QUARTZ-seq consistently performing best, followed by Smartseq2. Among droplet-based protocols, 10x Genomics yielded the highest number of detected genes in agreement with Mereu et al. Compared with the other single-cell methods, scMarathon-seq performs within the range of droplet-based methods. This is particularly notable given its current developmental stage. Since we had previously observed a tendency of uMRT to capture mitochondrial genes, we next compared mitochondrial read detection between scMarathon-seq and FS. Strikingly, for PBMCs, scMarathon-seq showed a lower overall fraction of mitochondrial reads than FS across all cell types (**Figure 6D**). This difference was most pronounced in CD14 cells (0.81 % ± 0.55 in scMarathon-seq, 6.67 % ± 1.29 for FS) and CD19 cells (3.81 % ± 2.31 and 10.07 % ± 2.25, respectively). For both RTs, CD3 cells showed the highest proportion of reads mapping to mitochondrial regions (7.55 % ± 3.02 for uMRT and 11.38 % ± 5.17). However, the distribution of the most highly expressed mitochondrial genes differed between uMRT and FS. Although scMarathon-seq captured fewer mitochondrial reads overall, these reads were distributed more evenly across mitochondrial genes, consistent with our observations in HeLa and K562 cells (**Figure 6E**). As previously observed, mitochondrial reads captured by FS predominantly cluster on the mitochondrial genes *MT-RNR2* and *MT-CO2*, whereas scMarathon-seq provides a more even profile of mitochondrial capture. More broadly, scMarathon-seq produced reads distributed across all chromosomes, although some differences in read abundance between uMRT and FS were observed for some chromosomes (**Supplementary Figure 6E**).

**Figure 6:**
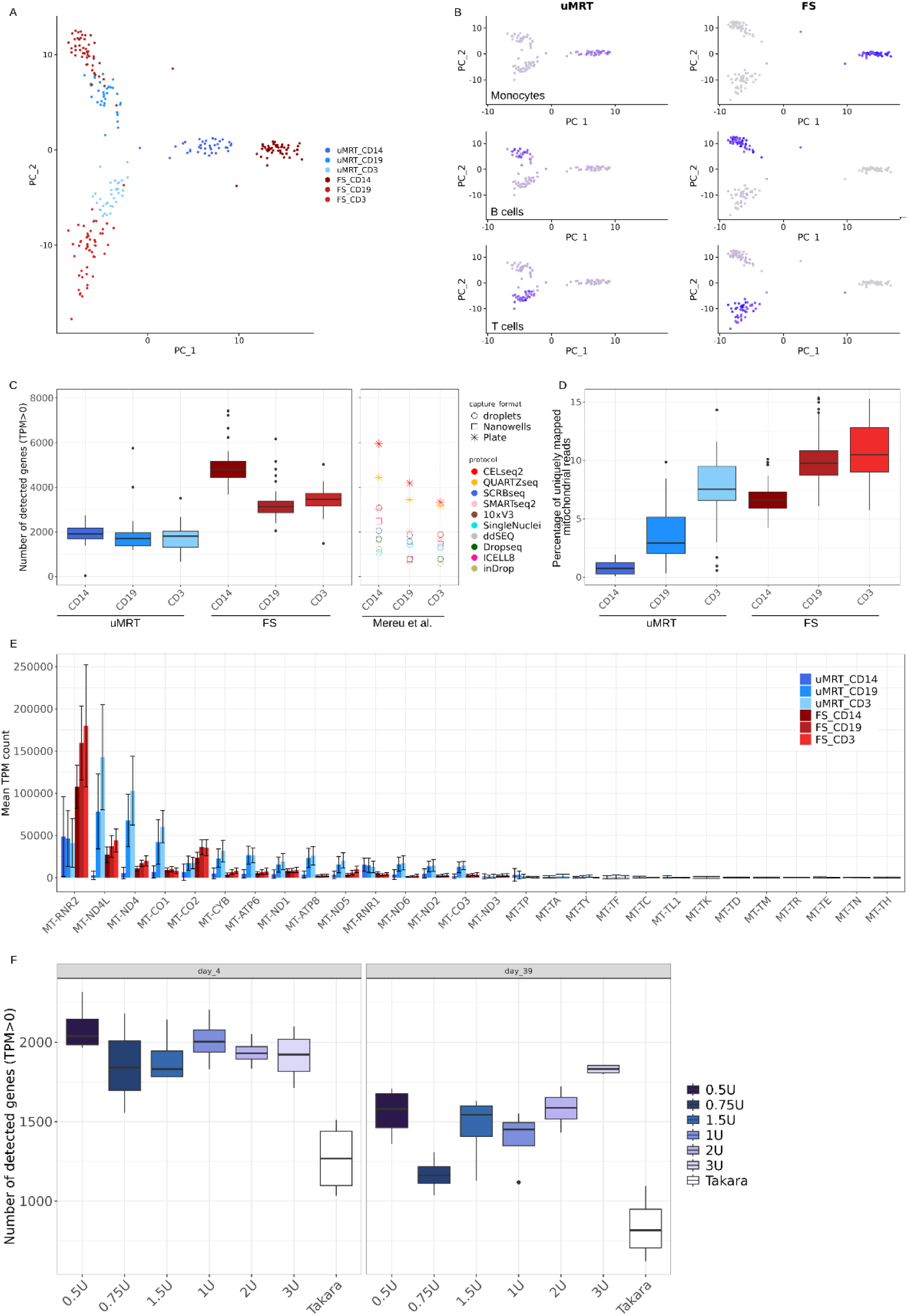
scMarathon-seq recovers PBMC transcriptomes and robustly recovers genes with SEQURNA after long-term storage. **(A)** Principal component analysis (PCA) of genes driving variability between PBMC cell types CD14, CD19 and CD3 between scMarathon-seq (n = 38, 33 and 25, respectively, blue shades) and FS (n=50) for each cell type, red shades), downsampled to 1 million reads. PCA was calculated on the 2,000 most variable genes, using 5 principal components. Mitochondrial and ribosomal genes were excluded from the analysis. **(B)** Projection of cell-type specific biological signatures onto PCA plots. Signatures were quantified as module scores based on marker genes for each cell type. Purple shades indicate signature strength relative to background gene expression. **(C)** Detected genes in PBMC cell types with scMarathon-seq and FS, compared with 11 single-cell/single-nuclei protocols benchmarked by Mereu et al. The left panel shows boxplots of detected genes number in PBMCs sequenced with scMarathon-seq and FS. The right panel shows median detected gene numbers from our reanalysis of PBMC cell-gene expression matrices from Mereu et al. Data points are shaped by capture format and coloured by protocol. MARSseq and Fluidigm-based samples were excluded. Genes were considered detected at TPM > 0. Boxplots show the median, interquartile range and whiskers show the full data range. **(D)** Boxlots depicting the percentage of reads falling within mitochondrial transcripts for CD14, CD19 and CD3 cells processed with scMarathon-seq (blue shades) and FS (red shades). Reads were mapped with STAR and only reads with a MAPQ score of 255 were included for the analysis. One CD3 cell of FS was removed as an outlier. **(E)** Distribution of gene expression of the highest-expressed mitochondrial genes between uMRT (blue shades) and FS (red shades). Gene expression is shown in TPM and mitochondrial gene annotation was extracted from GENCODE47. Error bars indicate the standard deviations between single cells. **(F)** Number of detected genes in CD4 cells after 4 days (left) and 39 days (right) of sample storage for SEQURNA concentrations ranging from 0.5-3 U/μl (n=4 per concentration, blue and purple shades) and the standard Takara inhibitor (n=4, white). Shading of the curves indicates the standard deviation between single cells. Genes were considered detected at TPM > 0.

**Supplementary Figure 6:**
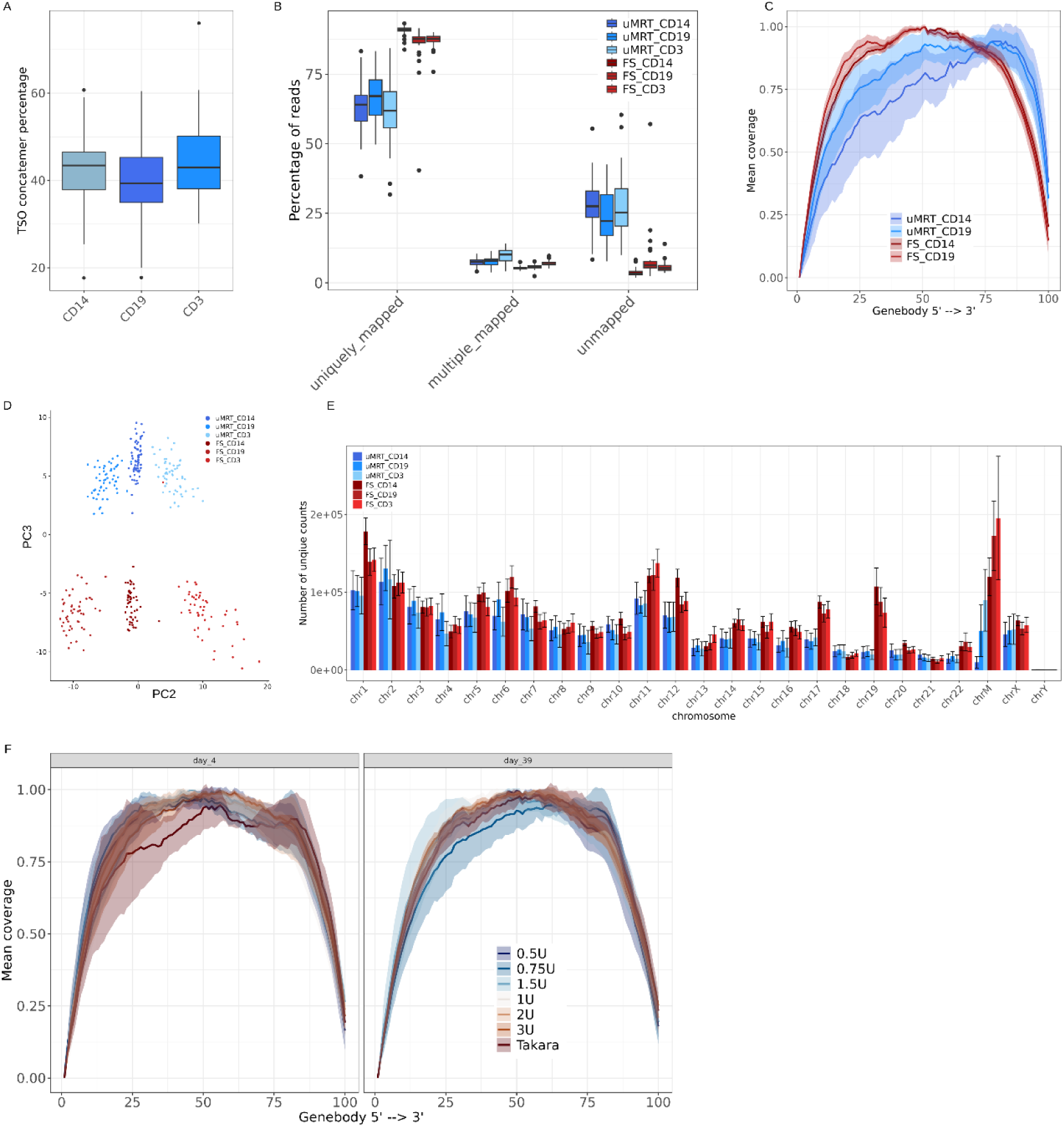
Quality metrics of PBMCs sequenced with scMarathon-seq and FS and SEQURNA-treated CD4 cells. **(A)** TSO concatemer rates in uMRT CD14, CD19 and CD3 cells (n=38, 33 and 35, respectively). Boxplots show the median with interquartile range and whiskers indicate the full data range. TSO concatemer rates were calculated by dividing the number of scMarathon-seq TSO concatemer containing reads by the total raw FASTQ reads for each sample. **(B)** Mapping statistics of uMRT (blue shades) and FS (red shades) for PBMC single cells. A read is considered to be uniquely mapped with a MAPQ score of 255. Reads were mapped with the STAR aligner using the GENCODE47 annotation. **(C)** Gene body coverage analysis of scMarathon-seq (blue shades) and FS (red shades) CD14 and CD19. Shown is the full length gene body coverage from 5’ to 3’Coverage binned into 100 intervals. **(D)** PCA of PC2 and PC3, with PC3 capturing technical variability between the uMRT and FS protocols. PCA was performed on the 2000 most variable genes using five principal components. Mitochondrial and ribosomal genes were excluded. **(E)** Distribution of uniquely mapping reads across each human chromosome for scMarathon-seq (blue shades) and FS (red shades). A read is considered uniquely mapped when its MAPQ score is 255. Error bars indicate standard deviation between single cells for each sample type. **(F)** Gene body coverage analysis of CD4 cells after 4 days (left) and 39 days (right) of sample storage for SEQURNA concentrations ranging from 0.5-3 U/μl (n=4 per concentration, blue and purple shades) and the standard Takara inhibitor (n=4, white). Shading of the curves indicates the standard deviations between single cells.

In parallel to assessing the recovery of PBMC transcriptome, we tested the efficacy of the RNAse inhibitor SEQURNA on CD4 cells. In contrast to conventional protein-based recombinant RNAse inhibitor (e.g. from Takara), SEQURNA is a synthetic, chemical RNA inhibitor that does not need reducing agents such as DTT and also poses no risk of cell-based contaminants from production (Noble et al., 2024). SEQURNA has been reported to detect 14-50 % more genes than Takara at 0.5-1 U/μl, at lower reagent cost and maintained long-term stability (Khven et al. 2026; Noble et al. 2024). We therefore sought to evaluate the performance of SEQURNA with scMarathon-seq after long-term storage and to determine the optimal conditions for our protocol. We compared the number of detected genes after 4 days of storage and 39 days of storage, across a concentration range from 0.5 U/μl to 3 U/μl and benchmarked these conditions against the standard Takara inhibitor. At both storage time points, Takara performed worst in terms of gene detection, with a median of 1,268 ± 229 after 4 days of storage, decreasing to 816 ± 206 after 39 days (**Figure 6F**). All SEQURNA tested concentrations yielded similar numbers of detected genes of around 2000 after 4 days of storage, with 0.5 U/μl showing the highest median number of detected genes at 2039 ± 260.84 and 1.5 U/μl showing the lowest with 1831 ± 171.33 genes. After 39 days of storage, gene detection decreased overall, most notably at 0.75 U/μl, with the number of detected genes dropping to 1162 ± 112.85. However, the 3U/μl showed the smallest decline, decreasing from 1922.5 ± 168.07 after day 4 to 1831.5 ± 29.03 after day 39. In addition to stabilizing gene detection, SEQURNA also supported even genebody coverage, also after 39 days of storage (**Supplementary Figure 6F**).

Taken together, our results show that scMarathon-seq can be applied to recover biological signatures, resolve cell type clustering and enable cell type annotation in PBMCs. scMarathon-seq already detects genes in the range of droplet-based protocols. Once more, we show again that scMarathon-seq captures a broader diversity of mitochondrial genes than FS, despite a lower overall fraction of mitochondrial reads. Furthermore, we show that combining scMarathon-seq with SEQURNA improves gene capture detection efficiency even after long periods of storage. Moreover, we tested multiple SEQURNA concentrations to determine the optimal working conditions for scMarathon-seq protocols. In summary, we demonstrate that, even in its current early developmental stage, uMRT recovers biologically meaningful information. We are certain that uMRT holds strong potential to expand the scope of transcriptomic profiling through iterative optimization.

### scMarathon-seq Enables Detection of Newly Synthesized RNA

We wanted next to evaluate the ability of uMRT to detect newly synthesized RNA using RNA metabolic labeling and nucleotide conversion-based approaches (Erhard et al. 2019). We applied metabolic labeling with 4-thiouridine (4sU) on HeLa cells at a concentration of 300 µM for 2 hours, and compared mismatch profiles across three sequencing technologies: Quant-seq with 10 ng extracted RNA input (bulk) as in (Herzog et al. 2017), Smart-seq technology from Takara Bio starting from a pool of 100 cells down to single-cell as in (Erhard et al. 2019), and scMarathon-seq from 10 ng extracted RNA (bulk) and a pool of 100 cells down to single-cell. Using matched labeled and unlabeled samples across a range of input sizes (bulk, mini-bulk, and single cell; see Methods), we were able to quantify the number of mismatches (Jürges et al. 2018). T→C mismatches, indicative of 4sU incorporation following nucleotide conversion, were enriched in labeled samples across all technologies and input sizes, while unlabeled controls showed low background frequencies (**Figure 7A; Supplementary Figure 7**). Background T→C mismatch frequencies were comparable between scMarathon-seq and Smart-seq, and most other mismatch frequencies were broadly similar across technologies at matched input sizes (**Supplementary Figure 7**).

**Figure 7:**
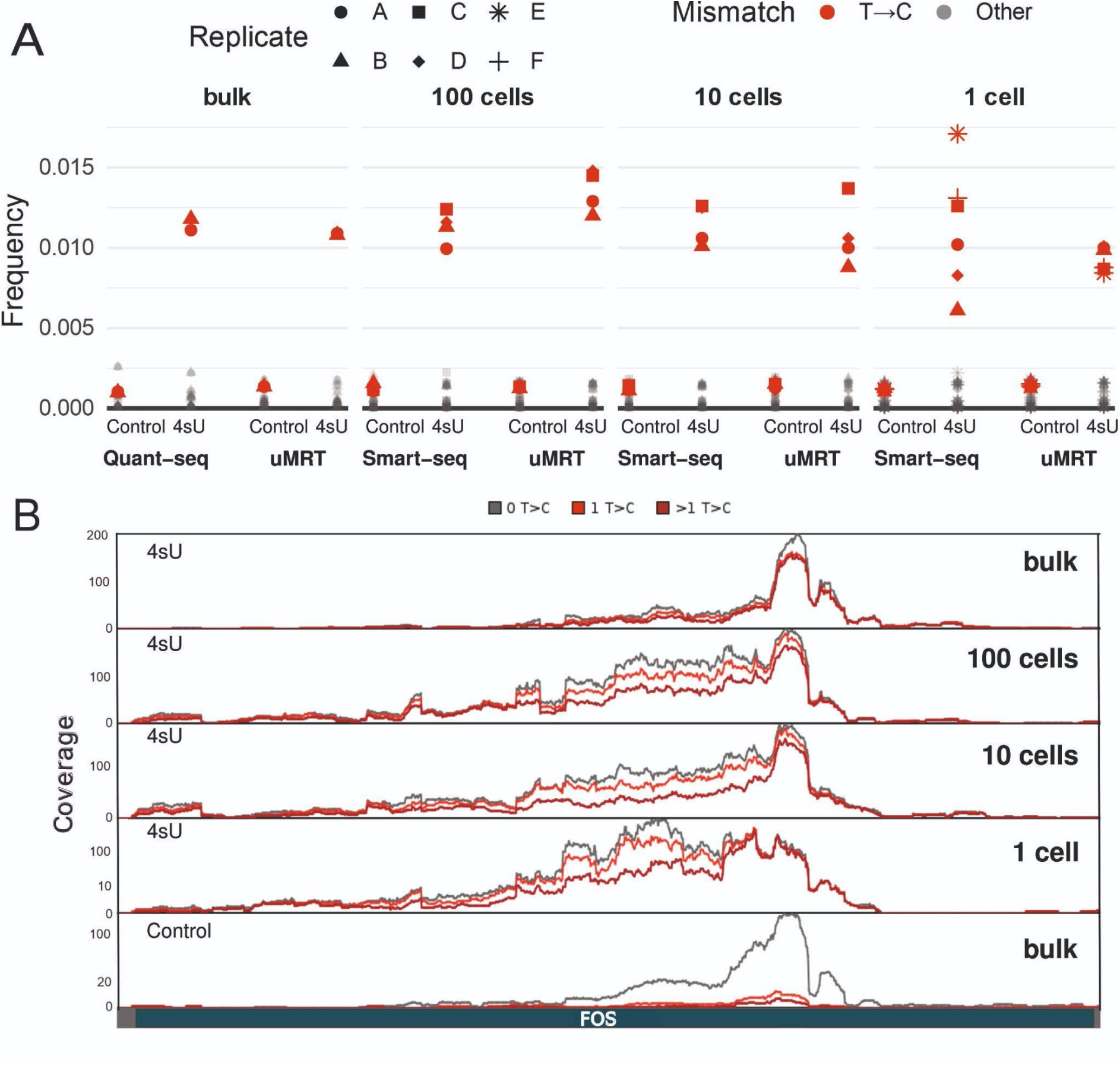
uMRT Detects 4sU-Induced T→C mismatches across bulk, mini-bulk and single-cell inputs. **(A)** Global nucleotide mismatch frequencies per sample, calculated using exonic sense reads only, in unlabeled controls and 4sU-labeled samples across sequencing technologies and input sizes. Bulk samples were analyzed using Quant-seq and uMRT; 100-cell, 10-cell, and single-cell samples were analyzed using Smart-seq and uMRT. T→C mismatches are shown in red, and all other mismatch types are shown in grey. Shapes indicate replicates. **(B)** Genome browser view of *FOS* coverage, highlighting T→C mismatches in uMRT samples. The upper four panels show 4sU-labeled samples across decreasing input sizes (bulk, 100 cells, 10 cells, and one cell). The bottom panel shows the unlabeled bulk control. Total coverage is shown in grey, coverage of reads containing at least one T→C mismatch is shown in red, and coverage of reads containing more than one T→C mismatch is shown in dark red.

**Supplementary Figure 7:**
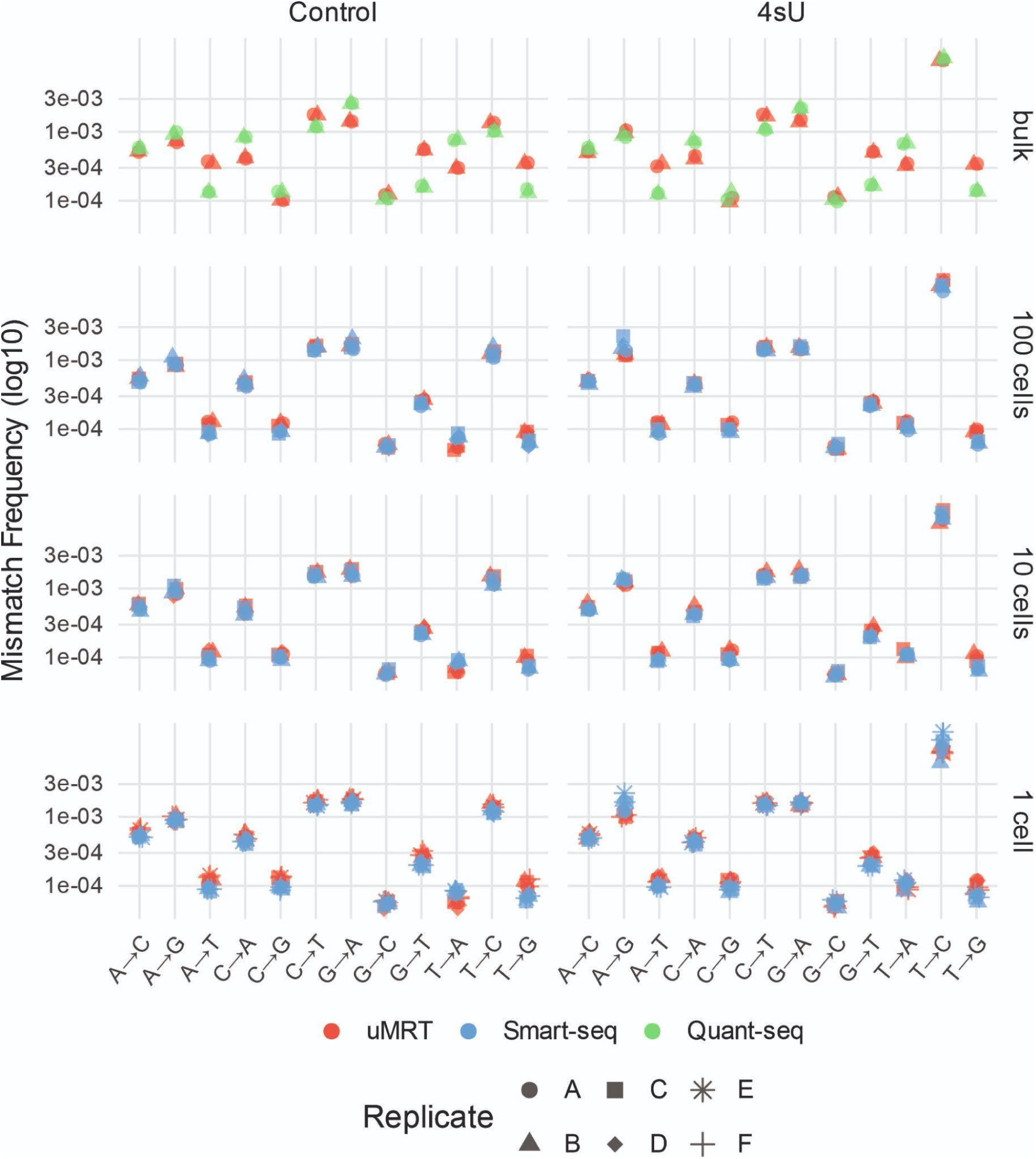
Global Mismatch Frequencies. Scatterplot of global nucleotide mismatch frequencies per sample, calculated using exonic sense reads only and colored by sequencing technology (Quant-seq, Smart-seq, uMRT). Panels show unlabeled controls and 4sU-labeled samples across input sizes (bulk, 100 cells, 10 cells, and one cell). Mismatch frequencies are shown on a logarithmic scale. Shapes indicate replicates.

These results demonstrate that uMRT retains a distinct labeling-associated T→C signal down to the single-cell level, supporting the differentiation of newly synthesized from pre-existing RNA. A genome browser view of the FOS locus further illustrates this capability: reads containing either one or multiple T→C mismatches are detectable in labeled uMRT samples across all input sizes, including single cells, whereas the unlabeled bulk control shows few such reads (**Figure 7B**). Taken together, these results demonstrate that uMRT can detect 4sU-induced T→C mismatches across a range of input sizes, indicating its suitability for metabolic RNA labeling and its potential for studying transcriptional dynamics.

## Discussion

The ability to copy RNA molecules into complementary DNA molecules using RTs has revolutionized the ability to profile RNA molecules (Stark et al. 2019). The success of MMLV-based RNA-seq workflows lies on their ability to perform template-switching, enabling versatile use across various protocol formats (e.g. plate-based, droplet-based), and facilitating easy-to-implement workflows with minimal experimental steps (Hahaut et al. 2022). This capability also enables picogram level of sensitivity, paving the way for performing scRNA-seq (Picelli et al. 2013). By systematically utilizing different experimental approaches through iterative experiments and 20 years of protocol optimization, some workflows now achieve exceptional transcript detection rates with over 10,000 transcripts per cell (Hagemann-Jensen et al. 2020, 2022; Hahaut et al. 2022) and reach down to femtogram level (Chen et al. 2026). Yet, the most significant developments have continuously focused on bringing scRNA-seq to a larger scale in order to reach organism-level profiling of mRNA (Regev et al. 2017), which comes at the expense of capturing the complete census of the single-cell transcriptomes (Quake 2021).

MMLV RT enzymes have inherent processivity limitations and are unable to unfold complex RNA molecules, which introduces bias in RNA-seq (Verwilt et al. 2023). Nonetheless, these artifacts have often been overlooked because no RT alternatives were available. Since the introduction of reverse transcriptases derived from group II intron retroelements as biotechnology tools (Mohr et al. 2013; Zhao et al. 2018), and especially the proof that these enzymes can perform RNA-seq (Nottingham et al. 2016; Guo et al. 2024), it is possible to envision developing a de-novo workflow for scRNA-seq.

Here, we have developed scMarathon-seq, a plate-based scRNA-seq protocol centered around UltraMarathonRT, an ultraprocessive RT able to perform template-switching (Guo et al. 2024). We systematically optimized lysis medium, the reverse transcription, and PCR parameters of the protocol to increase cDNA fragment length and the number of detected genes down to pg-level input (**Figure 1**). To benchmark scMarathon-seq performance, we chose FLASH-seq (FS) (Hahaut et al. 2022), a plate-based scRNA-seq protocol improved from the original Smart-seq2 (Picelli et al. 2013) and powered by Superscript IV, which has demonstrated the best gene-recovery capacity to date. Using two ENCODE cell lines, Hela and K562, perhaps not surprisingly, we could uncover major differences in the transcriptome captured by FS and scMarathon-seq using from ng level of RNA as input down to single-cell (**Figure 2, 3, 4**). Although both protocols retrieve the same number of genes, we see that more than 20-25% of the transcriptome is enzyme-dependent, with uMRT capturing longer genes. This difference is even more pronounced when we focus on lncRNAs, for which only one-third are profiled by both enzymes (**Supplementary Figure 2**). This ability to dissect gene composition is reflected in the data on the non-coding RNA *MALAT1*, which is an 8,7-kilobase-long RNA molecule that is rich in stable RNA secondary structures (Monroy-Eklund et al. 2023). Here we report a far more complex expression profile for *MALAT1* in uMRT-based data relative to MMLV-based data (**Figure 4H**).

An unexpected finding is that the scMarathon-seq drastically improves the profiling of mitochondrial poly(A) RNA in cell lines and primary cells compared to FS (**Figures 5 and 6**). The mitochondrial genome encodes 37 genes, giving rise to abundant polyadenylated mtRNA molecules (Jedynak-Slyvka et al. 2021). These can represent up to 50% of the total transcripts in a lysate mixture (Ludwig et al. 2019) and account for the most abundant transcripts (Boivin et al. 2018). Despite this abundance, mtRNA molecules are exceptionally challenging to profile, and their characterization has required complicated biochemical steps (Miller et al. 2022; Moran et al. 2024). This has caused significant technological issues, as mtRNA profiling is critical for successful lineage tracing. Given its ability to faithfully report mtRNA populations and isoforms, uMRT has the potential to pave the way for a new generation of assays that overcome the conventional burden of combining scATAC-seq and scRNA-seq (Miller et al. 2022; Ludwig et al. 2019).

Finally, we aimed to demonstrate that uMRT is compatible with time-resolved scRNA-seq using metabolic RNA labeling (**Figure 7**) (Erhard et al. 2022). Conceptually, we used 4-thiouridine (4sU) as a nucleotide analog that can be incorporated into newly synthesized RNA molecules, leading to T-C mismatches in the reads. We compared the Quant-seq workflow that was originally used to perform SLAM-seq (Herzog et al. 2017), Smart-seq using Takara kits at the single-cell level (Erhard et al. 2019), and the scMarathon-seq workflow. We established that 4sU incorporation can be effectively detected with uMRT down to the single-cell level. Altogether, it demonstrates that scMarathon-seq is not only amenable to unveiling the transcriptome at the single-cell level but also its dynamics.

## Methods

### Cell lines and single-cell sorting

ENCODE tier-one cell line K-562 (ATCC CCL-243) and tier-two cell line HeLa S3 (ATCC CCL-2) were purchased from ATCC. K562 cells were cultured in RPMI 1640 medium (Gibco) and HeLa S3 cells were cultured in DMEM (Gibco). Both types of culture media were supplemented with 10% v/v FBS (Fetal Bovine Serum, Gibco), 1% v/v Pencillin-Streptomycin (100 units/mL penicillin and 100 µg/mL streptomycin, Gibco), and 2 mM L-glutamine (200 mM, Gibco). Cells were passaged every second day and cultured at 37°C with 5% CO_2_ saturation. Mycoplasma absence from cell culture was regularly ensured using the Mycoplasma check service of Eurofins Genomics.

For harvesting, HeLa S3 cells were washed once in PBS (1x, Gibco) and detached with Trypsin-EDTA (0.25%, Gibco) for 5 min at 37°C. Then, trypsinization was stopped with ten volumes of medium before resuspending and collecting the cells. Since K562 cells grow in suspension the cells were just collected. Both cell lines were washed twice in PBS 1x, (Gibco) by centrifuging at 250 x g for 5 min, and resuspended in PBS (1x, Gibco) until further use.

Cells were resuspended in PBS (1x, Gibco) to reach a concentration of 1.10^6^ cells/100 µL. Live-dead cell assay was performed with 5 µL of Propidium Iodide Ready Flow Reagent (ThermoFisher) added to 100 µL cell suspension and incubated in the dark for 2 min on ice. The stained cell suspension was diluted three times in PBS (1x, Gibco) and passed through a 70-µm cell strainer (pluriSelect) before loading on the FACSAria Fusion Flow Cytometer (BD Biosciences). After setting up a gating strategy to exclude dead cells and doublets, single cells were sorted through a 100 µm nozzle into cooled, 48-well no-skirt PCR plates (Brand) prefilled with 2.4 µL lysis buffer (see below). Immediately after sorting, each plate was spun down in a plate centrifuge (Starlab) for 5 sec at maximum speed (500 x g) before transferring onto dry ice, and freezing at -80°C for storage. Plates containing single cells were thawed on ice for 3 min and spun down for 5 sec at 500 x g before use.

### PBMC isolation

Whole blood was drawn from a healthy donor into vials supplemented with anticoagulants (e.g EDTA). Within six hours after retrieval, the whole blood was diluted with one volume of room temperature 1x PBS (Gibco) and carefully pipetted on top of one volume of room temperature Histopaque®-1077 density gradient medium (Sigma-Aldrich) that was spun down in a fresh SepMate-50 tube (Stemcell Technologies) at 800 x g for 30 sec prior to use. The whole blood-Histopaque®-1077 mixture was centrifuged for 12 min at 1,200 x g at 22°C. After centrifugation, the yellow top layer was poured off into a new 50 ml Falcon tube without extensive transfer of red blood cells in the bottom layer, and centrifuged for 5 min at 800 x g at 22°C. The supernatant was aspirated, the pellet was resuspended in 1 mL of ACK buffer (Gibco) and incubated for 10 min on ice. The red blood cell lysis was quenched with 2 mL of ice-cold DPBS and centrifuged twice at 800 x g for 5 min at 4°C. For freezing, cells were resuspended in 2 mL of CryoStor® CS10 cell freezing medium (Sigma-Aldrich) in a concentration of 2 x 10^6^ cells per mL and stored at -80°C for short-term and liquid nitrogen for long-term storage.

Upon use, frozen cell vials were thawed in a water bath at 37°C for 2 min, and cells were transferred into fresh Falcon tubes. Then, the cells were diluted sequentially by incremental 1:1 volume additions of cell thawing medium at a speed of 1 mL per 5 sec and swirled with a 1-minute wait between each addition until 50 mL were completed. The tube was centrifuged at 400 x g for 5 min at room temperature. After carefully discarding the supernatant, the washing step was repeated with 25 mL of cell thawing medium. After the second wash, the cell pellet was resuspended in 1 mL of ice-cold FACS buffer (1x PBS + 1% FBS). Cell viability was examined using the Countess™ II automated cell counter (ThermoFisher). The expected viability was >90 % live cells.

### PBMC staining and single-cell sorting

After cell thawing, the cell suspension was centrifuged at 400 x g for 5 min at 4°C and gently resuspended with FACS buffer to a concentration of 1 x 10^6^ cells/100 µL. 100 µL of the cell suspension were transferred to a fresh 5 mL Eppendorf tube. To the sample, 5 µL of either anti-CD4-FITC (Biolegend), anti-CD8-FITC (Biolegend), anti-CD14-FITC antibody (Biolegend) or 4 µL of FITC isotype control (Biolegend) was added, mixed and incubated for 15 min on ice, protected from light. After the staining incubation, the cells were washed twice with 1 mL of FACS buffer and spun down at 400 x g for 5 min at 4°C. Finally, the cell pellet was resuspended in 100 µL FACS buffer and 5 µL Propidium Iodide Ready Flow Reagent (ThermoFisher) for 2 min in the dark. The cells were topped up to 1 mL with FACS buffer and kept in the dark and on ice until cell sorting. Prior to loading the sample into the FACSAria Fusion Flow Cytometer (BD Biosciences), the stained cell suspension was passed through a 70-µm cell strainer (pluriSelect) and carefully mixed by gentle vortexing. Proper gating ensured the exclusion of dead cells and doublets and selected for either CD4-positive, CD8-positive or CD14-positive single cells. Single cells were sorted through a 100 µm nozzle with the neutral density filter set to ND = 2.0 into cooled (4°C), 48-well no-skirt PCR plates prefilled with 2.4 µL lysis buffer. Immediately after sorting, each plate was sealed, spun down in a plate centrifuge (Starlab) for 5 sec at maximum speed (500 x g), transferred to dry ice, and frozen at -80°C until further processing.

### RNA extraction

For bulk input, approximately 1 million cells were collected for each cell line and washed twice in PBS (1x, Gibco). After the second wash, the supernatant was discarded and 300 µL RNA Lysis Buffer (Zymo Research) was added to the pellet. The pellet was immediately frozen on dry ice and stored at -80°C until further use. For RNA extraction, the Quick-RNA™ Microprep Kit (Zymo Research) was used as instructed by the manufacturer. In brief, the pellet was thawed at room temperature, resuspended with additional 300 µL absolute ethanol and transferred to a provided spin column before spinning at 16,000 x g for 30 sec. The column was then washed with 400 µL RNA Wash Buffer and treated with DNase for 15 min using a mixture of 5 µL DNase I and 35 µL DNA Digestion Buffer. Next, the sample was treated with 400 µL RNA Prep Buffer, followed by two consecutive washes with 700 µL and 400 µL RNA Wash Buffer. Finally, the RNA was eluted in 11 µL nuclease-free water.

### Total RNA extracts and synthetic RNA

Universal Human Reference RNA (UHRR) as a mix of human cell lines was purchased from ThermoFisher (#QS0639). Synthetic RNA from Spike-In RNA Variant Set 4 (SIRV-Set 4, Lexogen, #141.01) was spiked into the cell line samples (7.5 pg for bulk input and 0.225 pg for single cell input). All RNA samples were stored at stock concentrations in aliquots at -80°C for one-time use. Dilutions were prepared using nuclease-free water (Ambion) containing 0.4 units/µL recombinant RNase inhibitor (Takara Bio). RNA sample quality was monitored by the Bioanalyzer RNA Pico assay (Agilent).

### Marathon-seq

#### Cell lysis buffer and annealing

Plates for single cell-sorting (K562, HeLa S3 cells and PBMCs) were prefilled with 2.4 µL lysis buffer (0.15 µL Takara Lysis Buffer (10x, Takara Bio), 0.144 µL SEQURNA inhibitor (50 mass units/µL, Genovis), 0.72 µL betaine (5 M, Thermo Fisher), 0.6 µL uMRT_dT18 primer (5’-CCCTCTCTCTCTCTTTCCTCTCTCTTTTTTTTTTTTTTTTTT-3’, 5 µM, IDT), 0.3 µL dNTP mix (10 mM each, NEB), filled up with nuclease-free water (Ambion)) before sorting. For bulk input, 1 ng extracted RNA was added to the 2.4 µL lysis buffer. After thawing (single cells) and preparation (bulk), the samples were mixed on a ThermoMixer C (Eppendorf) at 4°C, 1,650 rpm for 30 sec, then spun down for 5 sec at 500 x g and incubated at 95°C for 15 sec and snap-cooled on ice for primer annealing. For experiments using UHRR, the lysis buffer contained 0.125 U/µL of recombinant RNAse inhibitor (Takara Bio) instead of SEQURNA.

#### Reverse transcription

To the primer-annealed RNA, an RT mix of 3 µL RT buffer (2x, RNAConnect), 0.3 µL Boost (RNAConnect), 0.1 µL uMRT enzyme (20 U/µL RNAConnect) and 0.2 µL nuclease-free water (Ambion) were added and mixed on a ThermoMixer C (Eppendorf) at 4°C, 1,650 rpm for 30 sec. Reverse transcription took place at 30°C for 30 min for scMarathon-seq.v2.

For scMarathon-seq.v1, the reaction used a homemade RT buffer (50 mM Tris-HCl, pH 8.3 at 25°C (Invitrogen), 200 mM KCl (Ambion), 4 mM MgCl_2_ (Ambion), 5 mM DTT (Sigma), 20 % glycerol (Promega)), additional 0.5 U/µL Recombinant RNase Inhibitor (Takara Bio) and was incubated at 42°C for 30 min.

#### Template-switching

Once reverse transcription was completed, 2.4 µL Template Switching Buffer (5x, RNAConnect), 1.2 µL TSO (5’*-CCCTCTCTCTCTCTTTCCTCTCTCTTTT-3’, 10 µM, RNAConnect, * modifications, TSO v2 / 2a are RNAConnect property), 1.2 µL dATP (10 mM, Carl Roth) and 1.2 µL uMRT enzyme (20 U/µL, RNAConnect) were added to a final reaction volume of 12 µL and mixed on a ThermoMixer C (Eppendorf) at 4°C, 1,650 rpm for 30 sec. For scMarathon-seq.v2, template switching occurred at 37°C for 15 min, followed by enzyme inactivation at 80°C for 5 min.

For scMarathon-seq.v1, the buffer was homemade (25 mM Tris-HCl pH 8.3 at 25°C (Invitrogen), 2 mM MgCl_2_ (Ambion), 2.5 mM DTT (Sigma), 10% PEG4000 (Sigma)) and incubation was performed at 30°C for 60 min, followed by inactivation at 80°C for 5 min.

#### PCR amplification

cDNA was amplified in a volume of 50 µL containing 5 µL AmpPCR primer (5’-CCCTCTCTCTCTCTTTCCTCTCTC-3’, 20 µM, IDT), 10 µL KAPA GC enhanced buffer (5x, Roche), 1.5 µL dNTP mix (10 mM each, Roche), 1 µL KAPA HiFi DNA polymerase (1 U/µL, Roche), filled up with 20.5 µL nuclease-free water (Ambion). After initial denaturation at 98°C for 2 min, the program was cycled 22-times for scMarathon.v2 and 25-times for scMarathon-seq.v1 at 98°C, 15 sec, 62°C, 30 sec and 72°C, 6 min. The final extension was performed at 72°C for 5 min. After PCR, the reaction was cleaned-up with AMPure XP beads (Beckman Coulter) in a 0.5x ratio and eluted in 12 µL TE buffer (Invitrogen). The PCR product concentrations were quantified with Qubit 1X dsDNA High Sensitivity (ThermoFisher), and length distribution was profiled on the Bioanalyzer using the High Sensitivity DNA Kit (Agilent).

#### Library preparation and sequencing

Sequencing libraries were prepared using the Nextera XT DNA Library Prep Kit (Illumina). In brief, 0.5 ng cDNA was tagmented using 2 µL Tagmentation DNA Buffer (TD) and 1 µL Amplicon Tagment Mix (ATM) in a thermocycler at 55°C for 15 min, with a hold step at 10°C. Immediately after the samples reach 10°C, 1 μL neutralizing buffer (NT) was added and incubated for 5 min at room temperature to strip off the Tn5 enzyme. The fragments were enriched with 3 µL Nextera PCR master mix (NPM), 1 µL Index 1 primer and 1 µL Index 2 primer in total reaction volume of 10 µL. The PCR program comprised denaturation at 72 °C for 3 min and at 95 °C for 30 s, followed by 12 cycles of denaturation at 95 °C for 10 s, annealing at 55 °C for 30 s, and extension at 72 °C for 30 s. A final extension was performed at 72 °C for 5 min. The libraries were cleaned up with AMPure XP beads in a 0.6x ratio (Beckman Coulter) and resuspended in 13 µL Resuspension buffer (RSB, Illumina). Library concentration was measured using the Qubit 1X dsDNA High Sensitivity or Quant-iT PicoGreen dsDNA Assay (ThermoFisher), and library size was visualized using the Bioanalyzer High Sensitivity DNA assay (Agilent). Finally, the libraries were pooled in equimolar concentrations and sequenced from both ends for 150 cycles on an Illumina NovaseqX for 7.5 million reads (single cells) or 20 million reads (bulk).

### FLASH-seq

FLASH-seq libraries were prepared as previously published (Hahaut et al. 2022).

#### Cell lysis buffer and annealing

Cells were sorted into plates prefilled with 1 µL lysis buffer consisting of 0.02 µL Triton X-100 (10%, Sigma), 0.24 µL dNTP mix (25 mM each, Carl Roth), 0.018 µL SMART dT30VN primers (5′Bio-AAGCAGTGGTATCAACGCAGAGTA CT30VN-3′, 100 µM, IDT), 0.03 µL recombinant RNase inhibitor (40 U/µL, Takara Bio), 0.012 µL DTT (100 mM, Invitrogen), 0.092 µL FS-TSO (5′Bio-AAGCAGTGGTAT CAACGCAGAGTACrGrGrG-3′, 100 µM, IDT), 0.09 µL dCTP (100 mM, Carl Roth), 0.2 µL betaine (5 M, ThermoFisher), 0.298 µL nuclease-free water (Ambion) and stored at -80°C. Upon use, the plates were thawed on ice, spun down for 5 sec at 500 x g and placed in a thermocycler at 72°C for 3 min.

#### RT-PCR

For RT-PCR, a 4 µL mix of 0.238 µL DTT (100 mM, Invitrogen), 0.046 µL MgCl2 (1 M, Ambion), 0.8 µL betaine (5 M, ThermoFisher), 0.096 µL recombinant RNase inhibitor (40 U/µL, Takara Bio), 0.050 µL SuperScript IV (200 U/µL, Invitrogen), 2.5 µL KAPA HiFi HotStart ReadyMix (2x, Roche), 0.270 µL nuclease-free water (Ambion) was added and placed in a thermocycler to run the following program: 50°C for 60 min, 98°C for 3 min, then cycled for 18 times at 98°C for 20 sec, 67°C for 20 sec and 72°C for 6 min. For clean-up, 10 µL of nuclease-free water (Ambion) was added to the cDNA libraries before mixing with 12 µL of AMPure XP beads (0.8x ratio, Beckman Coulter). The bead suspension was incubated for 5 min at room temperature and placed on a magnet for another 5 min. Once the solution became clear, the supernatant was discarded, and 15 µL nuclease-free water (Ambion) was added to resuspend the beads. The cDNA was incubated for 2 min at room temperature and placed on a magnet for another 2 min. 14 µL of the cleared solution was transferred to a fresh plate. cDNA was quantified using Qubit 1X dsDNA High Sensitivity (ThermoFisher), and profiled with the Bioanalyzer High Sensitivity DNA assay (Agilent).

#### Library preparation and sequencing

For library preparation, the Nextera XT DNA Library Prep Kit (Illumina) was used. Briefly, 1 µL cDNA was diluted to 0.15 ng/µL. From this dilution, 1 µL was mixed with 3 µL tagmentation mix (0.27 µL Amplicon Tagment Mix (ATM), 2.18 µL Tagmentation DNA Buffer (TD), 0.55 µL nuclease-free water (Ambion)) and incubated at 55°C for 8 min, followed by a hold at 4°C. Upon completion, 1 µl neutralization buffer (NT) was immediately added, mixed and incubated for 5 min at room temperature. Next, 2 µL premixed custom N7xx and S5xx index adapters (5 µM each, IDT) and 3 µL Nextera PCR master mix (NPM) were added and fragments were enriched using the following program: 72°C for 3 min, 95°C for 30 sec, 14 cycles of 95°C for 10 sec, 55°C for 30 sec, 72°C for 30 sec, hold at 15°C. After enrichment, 5 µL of each library were pooled into a single 1.5 mL Eppendorf tube, mixed with a 0.8x ratio of AMPure beads and incubated for 5 min. Then, the tube was placed on a magnet for another 5 min. Once the solution is cleared, the supernatant is discarded and the beads are washed with 1 mL 80% v/v ethanol for 30 sec. After removal of the ethanol and any residuals, the bead pellet was dried for 2 min until small cracks appeared. Then, the beads were resuspended in 50 µL nuclease-free water (Ambion) and incubated for 2 min for elution. Once again, the sample was placed on a magnet for 2 min and 49 µL of the clear solution was transferred to a new tube. The library quality was assessed with the Qubit 1X dsDNA High Sensitivity assay (ThermoFisher), and profiled with the Bioanalyzer High Sensitivity DNA assay (Agilent). Libraries were sequenced from both ends for 150 cycles on an Illumina NovaseqX sequencer at a depth of 1.67 million reads for single cells and 10 million reads for bulk input.

### scSLAM-seq

#### 4-thiouridine labeling

One day prior to the experiment, 5.10^5^ HeLa S3 cells were seeded in a T25 flask to reach confluency the next day. On the day of the experiment, 30 µL 4-thiouridine (50 mM, Biosynth) was added to 5 mL DMEM medium (Gibco) and incubated for 2 hours. Then, cells were harvested by detaching with 0.5 mL Trypsin-EDTA (0.25%, Gibco) for 5 min. Trypsin treatment was stopped with 4.5 mL DMEM medium and cells were washed twice in PBS (1x, Gibco) by centrifuging at 250 x g for 5 min. Cell sorting (single cells, 10 cells, and 100 cells) on plates was performed as described above.

#### Alkylation of 4-thiouridine

Wells containing single cells in 2.4 µL lysis buffer for Marathon-seq (0.15 µL Lysis Buffer (10x, Takara Bio), 0.144 µL SEQURNA inhibitor (50 mass units/µL, Genovis)) or 4 µL lysis buffer for SMART-seq (0.4 µL Lysis Buffer (10x, Takara Bio), 0.2 µL recombinant RNase inhibitor (40 U/µL, Takara Bio)) were filled to 4 µL with nuclease-free water (Ambion). 0.4 µL PBS pH 7.4 (10x, Gibco) were added to all wells. Then, 4.4 µL Pierce™ Iodoacetamide (20 mM, ThermoFisher) resuspended in DMSO (Sigma) was added to all wells and briefly centrifuged. Alkylation took place during an incubation of 5 min at 50°C in a thermocycler. The reaction was quenched with 1.3 µL DTT (0.1 M, ThermoFisher) for 5 min at room temperature.

#### In-well clean-up step

10.1 µL of RNA XP beads (1:1 ratio, Beckman Coulter) were added to the reaction, mixed well and incubated for 10 min at room temperature. Then, the samples were placed on a magnet for 4-5 min until the supernatant became clear. The supernatant was discarded, and the beads were washed twice for 30 sec with 50 µL of 80% v/v ethanol. After the beads were air-dried for a maximum 2 min, the RNA was resuspended and incubated for 3 min at room temperature in elution buffer consisting of the components for annealing depending on the protocol (Marathon-seq or SMART-seq). For elution, the samples were centrifuged at 500 x g for 2x 30 sec pulses until the bead pellets formed at the bottom of the wells. From that point on, samples were centrifuged at 500 x g for 2x 30 sec pulses before every incubation step to avoid bead interference with downstream reactions.

#### Marathon-seq

The RNA beads were resuspended and incubated for 5 min at room temperature in 1.68 µL elution buffer consisting of 0.144 µL SEQURNA inhibitor (50 mass units/µL, Genovis), 0.6 µL uMRT_dT18 primer (5’-CCCTCTCTCTCTCTTTCCTCTCTCTTTTT TTTTTTTTTTTTT-3’, 5 µM, IDT), 0.3 µL dNTP mix (10 mM each, NEB) and 0.64 µL nuclease-free water (Ambion). After centrifuging at 500 x g for 2x 30 sec pulses, 0.72 µL betaine (5 M, ThermoFisher) was added. The remaining steps were performed as described above. The samples were sequenced paired-end for 150 cycles at a depth of 8 million reads on an Illumina NovaseqX sequencer.

#### SMART-seq

All reagents were provided by the SMART-Seq® v4 Ultra® Low Input RNA Kit (Takara Bio). The RNA beads were resuspended and incubated for 5 min at room temperature in 4 µL elution buffer consisting of 0.2 µL recombinant RNase inhibitor (40 U/µL, Takara Bio), 0.5 µL CDS primer, and 3.3 µL nuclease-water. After centrifuging at 500 x g for 2x 30 sec pulses, the samples were annealed at 72°C for 3 min in a pre-heated thermocycler, followed by a cool-down to 4°C for 3 min. Reverse transcription readily followed by adding 1 µL 5X Ultra-low first strand buffer, 0.25 µL SMART-seq v4 Oligo, 0.125 µL RNase inhibitor, 0.5 µL SMARTScribe Reverse Transcriptase and incubating at 42°C for 90 min, then at 70°C for 10 min. Second strand synthesis and amplification was performed in 6.25 µL 2X SeqAmp PCR Buffer, 0.25 µL PCR Primer II A, 0.25 µL SeqAmp DNA Polymerase and 0.75 µL nuclease free-water at 95°C for 1 min, 21 cycles at 98°C for 10 s, 65°C for 30 sec, 68°C for 3 min and 72°C for 10 min for final extension. cDNA libraries were cleaned-up by adding 12.5 µL AMPure XP beads (Beckman Coulter) supplemented with 0.25 µL 1x Lysis buffer (Takara Bio) and incubating for 8 min at room temperature. Next, the beads were washed twice for 30 sec in 100 µL 80% v/v ethanol, before air-drying and eluting in 17 µL elution buffer (Takara Bio). cDNA was quantified using Qubit 1X dsDNA High Sensitivity (ThermoFisher), and profiled with the Bioanalyzer High Sensitivity DNA assay (Agilent). Libraries were prepared using the Nextera XT DNA Library Prep Kit (Illumina) as described for Marathon-seq. All samples were sequenced paired-end for 150 cycles on an Illumina NovaseqX sequencer at a depth of 8 million reads.

### Data analysis of RNA-seq and scRNA-seq

#### Preprocessing and main data analysis scripts for bulk and single-cell RNA-seq

TSO concatemers of raw paired-end and single-end sequencing reads were counted with a custom bash script. A TSO concatemer was classified as any read containing more than one occurrence of the TSO sequence or its reverse complement. Raw sequencing reads were trimmed with fastp (v0.23.2) to remove sequencing adapters, TSO sequences, and G polymers. The trimmed reads were mapped with STAR (v2.7.11b) to the human reference genome GRCh38 (GENCODE 47, GRCh38 primary assembly) and subsequently uniquely mapped reads were quantified for each gene using featureCounts (v2.0.6) in paired-end mode without allowing overlap using the following parameters : -p –countReadPairs -s 0 -t exon -g gene_id. To normalize for gene length, the resulting raw counts per gene were transformed into TPM using a custom R script. Pairwise correlations between replicates and samples were calculated with log10(TPM+1). Genes were considered as ‘detected’ with at least one count in one replicate and sample and considered as expressed with TPM ≥ 1, respectively. Gene body coverage was calculated using RSeQC (v5.0.1). ‘Percentages of mitochondrial reads’ were calculated by dividing the number of unique mitochondrial mapping reads by the sum of total uniquely mapping reads (MAPQ 255) per sample. To schematize the mitochondrial chromosome, the package karyoploteR (1.24.0) was used and the corresponding mitochondrial genes symbols along with their genomic positions were added by using the R package biomaRt (v2.54.0). To visualize read coverage, samtools (v.1.13) depth was used to obtain the read depth at each base for needed genomic ranges. Gene models were created with the R package ggtranscript (1.0.0) based on the GENOCDE 47 annotation file. Union exon numbers and unified intronic length were determined with the R packages GenomicAlignments (v1.34.9), GenomicFeatures(v.1.50.2) and GenomicRanges (v.1.50.0). Gene lengths and corresponding biotypes were extracted from the geneInfo.tab file obtained after mapping with STAR. Furthermore, the RSeQC (v5.0.1) software package was used to examine the distribution of reads across different genomic features (CDS, 5′UTR, 3′UTR, introns, TSS_up_10kb, TES_down_10kb). In order to investigate the distribution of reads in more detail, the underlying RSeQC script read_distribution.py was modified to extract each read and its corresponding read information (genomic start and end position,CIGAR length, read sequence) of each genomic element. Downstream analysis was conducted using R (v.4.2.3) and Rstudio (v2022.07.2). For sequencing saturation analysis and to account for the effect of different sequencing depths, reads were downsampled with seqtk (v1.4). Portions of the data analysis, code development, spelling checks and text formatting were assisted using ChatGPT (GPT-5.5, OpenAI).

#### Analysis of publicly available K562 and HeLa single-cell datasets

uMRT performance was compared to two publicly available 10x Genomics datasets. For K562 cells, the feature/cell matrix from the dataset 10k_K562_singleplex, consisting of K562 cells Fixed RNA profiling from ATCC (CRL-3344) by Cell Ranger v7.0.0, 10x Genomics (2022, May 13) was analyzed. For HeLa cells, a publicly available UMI count matrix of HeLa S3 scRNA-seq was downloaded from the National Center for Biotechnology Information (NCBI) Gene Expression Omnibus (GSE138814). Both datasets were processed with Seurat’s (v5.2.1) standard workflow. For HeLa, only cells containing more than 200 and less than 8000 genes and less than 10 % of mitochondrial genes were retained for downstream analysis. For K562, cells were retained for downstream analysis if they comprised more 200 and less than 9000 genes with a percentage of less than 10 % of mitochondrial genes. Cell counts were normalized and scaled with the default settings of dividing each cell by its total expression, multiplying it by a scale factor of 10e6 and log-transforming the result. Highly variable features were determined with the default parameters of Seurat’s FindHighlyVariableFeatures() function, utilizing the 2000 features showing the highest variability of the mean-variance relationship after using local polynomial regression (loess). The number of detected genes was then determined by calculating the median number of genes per cell.

#### Dimensionality reduction of HeLa and K562 single-cell data

PCA was computed using Seurat’s (v5.2.1) RunPCA() function on HeLa and K562 raw count matrices after filtering out ribosomal and mitochondrial reads. The counts were normalized with Seurat’s default parameters and the 2000 most highly variable features were computed. After scaling the data, PCA was run on the highly variable genes with a number of 10 principal components.

#### Data analysis of single-cell PBMC data with Seurat

PBMC single-cell data was processed with Seurat’s (v5.2.1) standard workflow. Raw count matrices were normalized with default settings. The top 2000 most highly variable features were computed and the count matrix was scaled with default parameters. Linear dimensionality reduction was performed with PCA on highly variable features. Module scores for cell type identities were computed with Seurat’s AddModuleScore(). For the CD14 signature, the genes *CD14*, *VCAN*, *S100A8*, *S100A9*, *LYZ* and *FCN1* were used. CD19 signature was defined with the genes *MS4A1*, *CD19*, *CD79A* and *BANK1*. Lastly, the genes *CD3D*, *CD3E*, *TRAC*, *IL7R* and *CCR7* were utilized to compute the CD3 module score.

#### Data analysis of the benchmarking dataset of Mereu et al., 2020

Human cell-gene expression matrices were downloaded from the super series GSE133549 from the NCBI Gene Expression Omnibus as uploaded by (Mereu et al. 2020). Preprocessing of the provided cell-gene expression matrices for each protocol was executed following the supplementary R scripts of Mereu et al. To ensure the analysis of the same cells as in Mereu et al., cells with more than 25 % of mitochondrial reads were discarded, as well as cells containing fewer genes than the determined lowest 5th percentile of detected genes per protocol. Downstream analysis was conducted with Seurat (v5.2.1). Normalization of gene expression was performed by dividing each cell by its total expression, multiplying it by a scale factor of 10e4 (instead of the standard scale factor of 10e6) and log-transforming the result. For simple cell type annotation, highly variable features were computed with Seurat’s default FindVariableFeatures() function, utilizing the 2000 features showing the highest variability. Genes were scaled with the standard Seurat gene scaling procedure. Library sizes and mitochondrial genes were regressed out. Linear dimension reduction was performed by Principal Component Analysis (PCA) with Seurat’s RunPCA() function on the determined highly variable features. A shared-nearest-neighbour graph on the first 8 principal components was constructed in euclidean distance with Seurat’s FindNeighbours followed by clustering with the Louvain algorithm at a resolution of 0.4. To visualize cell clustering and aid in cell type annotation, non-linear dimensional reductions with t-SNE and UMAP were applied. Marker genes denoting cell clusters were identified by applying standard Wilcoxon rank-sum test with Seurat’s FindAllMarkers() function. For the scope of our comparison, we excluded HEK293T cells and included only PBMCs, specifically CD14, CD19 and CD3 cells. Markers utilized for annotating CD14 monocytes were *CD14*, *FCN1*, *LYZ*, *S100A8*, *S100A9*, *VCAN.* CD19 B cells were identified with the marker genes *BANK1*, *CD19*, *CD79A* and *MS4A1*. For CD3 T cells, we utilized the marker genes *CCR7*, *CD3D*, *CD3E*, *IL7R* and *TRAC*. MARS-seq and Fluidigm protocols were excluded from the comparison as cell type clustering proved to be too ambiguous to confidently assign cellular identities.

### scSLAM-seq analysis

Quant-seq, Smart-seq, and uMRT sequencing data were processed using GEDI (v1.0.6f; Erhard et al., 2018) and GRAND3 (v3.0.9). Smart-seq and uMRT reads were processed in paired-end mode, whereas Quant-seq reads were processed in single-end mode. Adapter sequences were removed from Smart-seq and uMRT reads using Cutadapt (v3.5). The sequences CCCTCTCTCTCTCTTTCCTCTCTCTTTT and AAAAGAGAGAGGAAAGAGAGAGAGAGGG were supplied for read 1 and read 2, respectively, and CTGTCTCTTATACACATCT was supplied for both mates. Quality trimming was performed using --nextseq-trim=20.

Reads were aligned to rRNA (U13369.1) and Mycoplasma databases with Bowtie2 (v2.3.0), followed by alignment of the remaining reads to the human GRCh38 primary assembly using STAR (v2.7.10b), with GENCODE release 47 gene annotation and the following parameters: --outFilterMismatchNmax 20, --outFilterScoreMinOverLread 0.4, -- outFilterMatchNminOverLread 0.4, and --alignEndsType Extend5pOfReads12. Aligned reads were converted and merged using GEDI Bam2CIT and MergeCIT. GRAND3 was then used (- auto -profile) to generate gene-level read counts, new-to-total RNA ratios (NTRs), and global nucleotide mismatch statistics (Jürges et al. 2018). Genome browser tracks were generated using GEDI Display, showing total coverage and coverage of reads containing at least one or more than one T→C mismatch. For mismatch frequency plots, GRAND3 output was restricted to the “Exonic (h.gencode47)” category and the “Sense” subread category. Plots were generated in R using ggplot2.

## Data and code availability

RNA-seq data generated in this study and data scripts that support the analysis in this manuscript are available upon request. Publicly available 10x Genomics for HeLa cells was downloaded from the National Center for Biotechnology Information (NCBI) Gene Expression Omnibus (GSE138814). Publicly available 10x Genomics for K562 cells was downloaded from 10x Genomics 10k_K562_singleplex dataset, consisting of K562 cells Fixed RNA profiling from ATCC (CRL-3344) by Cell Ranger v7.0.0, 10x Genomics (2022, May 13). Publically available processed gene expression files of PBMC as prepared by (Mereu et al. 2020) were downloaded from the NCBI Gene Expression Omnibus (Series GSE133549).

## Acknowledgments

We thank Brenton Graveley for advice and discussions on the experiments. This work was supported by the National Institutes of Health (1R01HG011868). A.M.L., F.E. and A.-E.S. thank the Deutsche Forschungsgemeinschaft (DFG) Collaborative Research Center 1583 DECIDE (Project Z2, DFG project number: 492620490) and Cluster for Nucleic Acid Sciences and Technologies – NUCLEATE (Project-ID 533767322 – EXC 3113/1) to A.-E.S.. A.E.S. and S.I.B. thank the Agence Nationale pour la Recherche (ANR)-DFG support MAID (#530098272). A.-E.S. thank the Helmholtz Association for funding Helmholtz Exzellenznetzwerk Phase 2 (EXNET-02-31). A.G. thanks the Munich School of Data Science (MUDS). C.-L.C. thanks the HIRI Graduate Program “RNA and Infection”.

## Author Contributions Statement

Conceptualization, A.M.P. and A.-E.S.; Methodology, C.-L.C., L.T.G. and S.B.; Software/data analysis, A.G., T.R., and F.E.; supervision, A.M.P. and A.-E.S.; writing – original draft, A.-E.S., C.-L.C. and A.G.; writing – review & editing, All co-authors.

---

**Supplementary Table 2:**
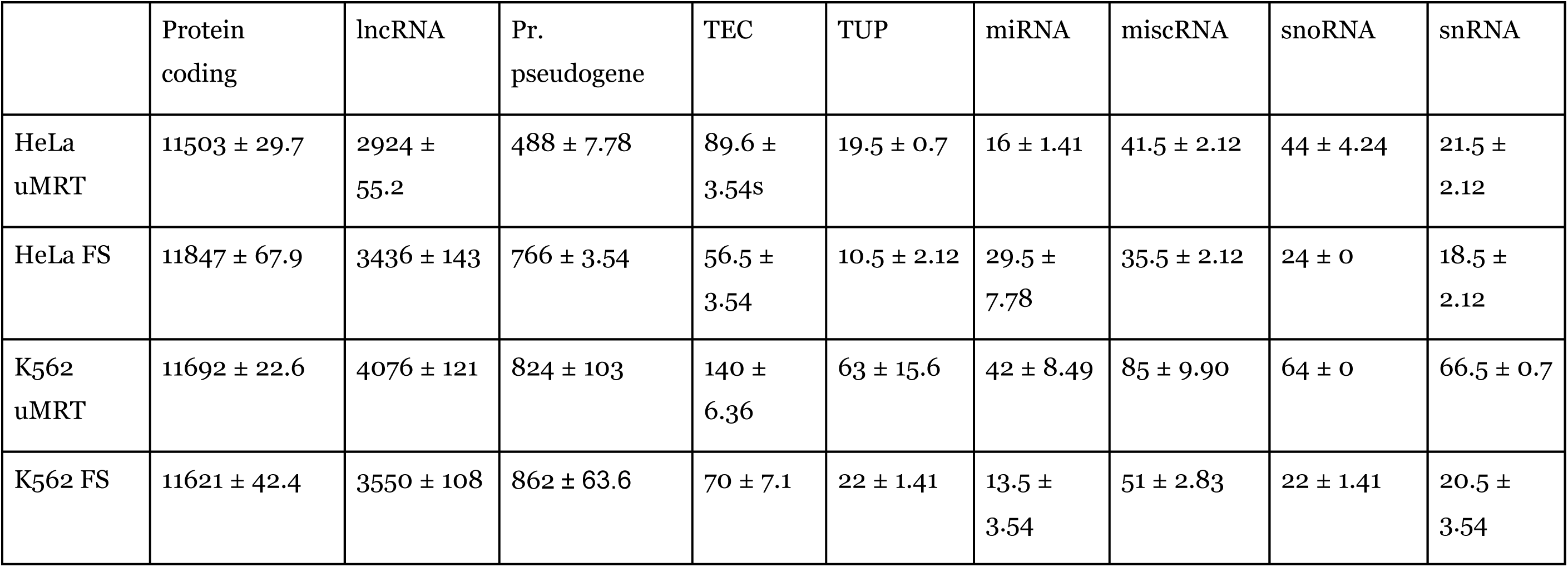
Detailed biotype distribution for Bulk RNA-seq of HeLa cells and K562 cells between scMarathon-seq and FLASH-seq. Exact percentages of GENCODE47 described biotypes with their respective standard deviation.

|  | Protein coding | lncRNA | Pr. pseudogene | TEC | TUP | miRNA | miscRNA | snoRNA | snRNA |
| --- | --- | --- | --- | --- | --- | --- | --- | --- | --- |
| HeLa uMRT | 11503 ± 29.7 | 2924 ± 55.2 | 488 ± 7.78 | 89.6 ± 3.54s | 19.5 ± 0.7 | 16 ± 1.41 | 41.5 ± 2.12 | 44 ± 4.24 | 21.5 ± 2.12 |
| HeLa FS | 11847 ± 67.9 | 3436 ± 143 | 766 ± 3.54 | 56.5 ± 3.54 | 10.5 ± 2.12 | 29.5 ± 7.78 | 35.5 ± 2.12 | 24 ± 0 | 18.5 ± 2.12 |
| K562 uMRT | 11692 ± 22.6 | 4076 ± 121 | 824 ± 103 | 140 ± 6.36 | 63 ± 15.6 | 42 ± 8.49 | 85 ± 9.90 | 64 ± 0 | 66.5 ± 0.7 |
| K562 FS | 11621 ± 42.4 | 3550 ± 108 | 862 ± 63.6 | 70 ± 7.1 | 22 ± 1.41 | 13.5 ± 3.54 | 51 ± 2.83 | 22 ± 1.41 | 20.5 ± 3.54 |

**Supplementary Table 3:**
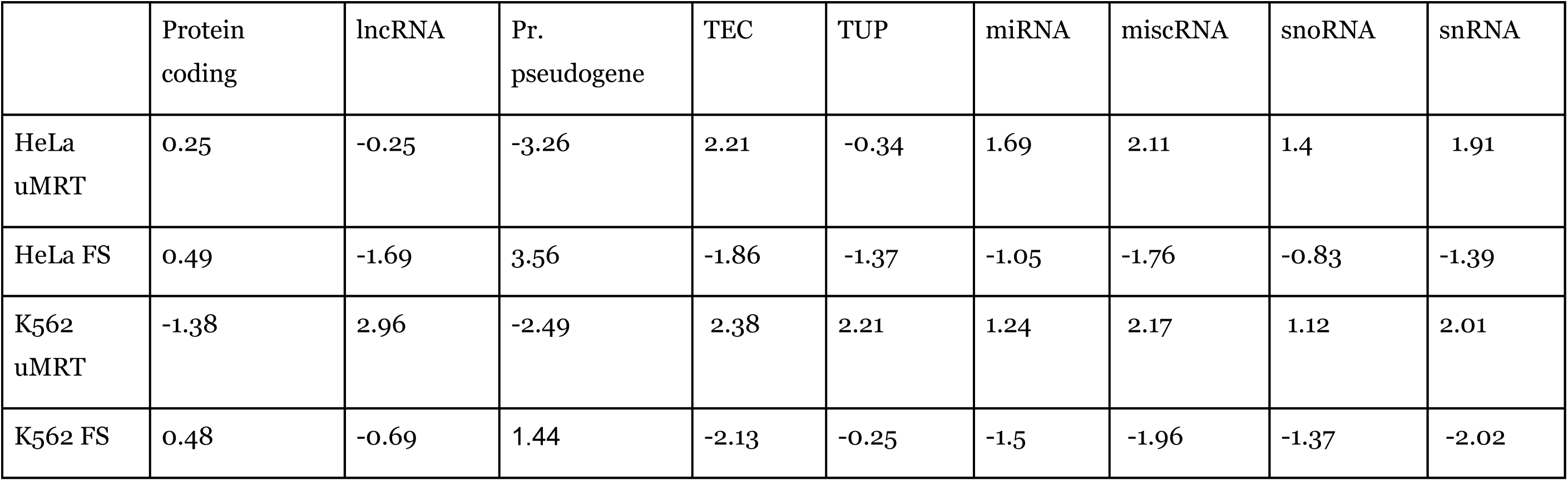
Detailed Pearson residuals of the biotype distribution for single-cell RNA-seq of HeLa cells and K562 cells between uMRT and Flash-seq. Pearson residuals were extracted from a Chi-squared test of independence between both protocols and cell types.

|  | Protein coding | lncRNA | Pr. pseudogene | TEC | TUP | miRNA | miscRNA | snoRNA | snRNA |
| --- | --- | --- | --- | --- | --- | --- | --- | --- | --- |
| HeLa uMRT | 0.25 | -0.25 | -3.26 | 2.21 | -0.34 | 1.69 | 2.11 | 1.4 | 1.91 |
| HeLa FS | 0.49 | -1.69 | 3.56 | -1.86 | -1.37 | -1.05 | -1.76 | -0.83 | -1.39 |
| K562 uMRT | -1.38 | 2.96 | -2.49 | 2.38 | 2.21 | 1.24 | 2.17 | 1.12 | 2.01 |
| K562 FS | 0.48 | -0.69 | 1.44 | -2.13 | -0.25 | -1.5 | -1.96 | -1.37 | -2.02 |

**Supplementary Table 4:** Detailed Pearson residuals of the biotype distribution for single-cell RNA-seq of HeLa cells and K562 cells between of uMRT-only and FLASHseq-only captured genes. Pearson residuals were extracted from a Chi-squared test of independence between both protocols and cell types.

|  | Protein coding | lncRNA | Pr. pseudogene | TEC | TUP | miRNA | snoRNA | snRNA |
| --- | --- | --- | --- | --- | --- | --- | --- | --- |
| HeLa uMRT | -7.86 | 7.94 | -0.54 | 7.29 | 2.1 | 1.11 | 3.41 | 1.11 |
| HeLa FS | 3.42 | -3.46 | 0.23 | -3.12 | -0.91 | -0.48 | -1.49 | -0.48 |
| K562 uMRT | -10.94 | 11.16 | -0.05 | 7.76 | 5.46 | 2.84 | 2.73 | 4.53 |
| K562 FS | 5.27 | -5.38 | 0.03 | -3.74 | -2.63 | -1.37 | -1.31 | -2.18 |

